# Host Cell Wall Polysaccharides Modulate Nanosynbacteria Recognition and Attachment

**DOI:** 10.64898/2026.10.06.756255

**Authors:** Alex S. Grossman, João Gabriel S. Souza, Lei Lei, Kamilla Anochshenko, Mengying Xia, Yaxi Wang, Jean-Lou Reyre, Elisabeth C. Lowe, Xuesong He, Patrick J. Moynihan, Joseph D. Mougous, Batbileg Bor

## Abstract

Ultrasmall episymbiotic bacteria grow on the surface of their obligate bacterial hosts.

These episymbionts have a restricted host range and likely rely on host-derived receptor molecules to identify and associate with cognate hosts. Nanosynbacteria from the oral microbiome, like *Nanosynbacter lyticus*, utilize a pair of distinct type 4 pili (T4P) for host surface interactions, T4P-1 and T4P-2. In this study we identify the adhesin components of both T4P, demonstrate that these adhesins localize to the cell wall of the cognate host Actinobacterium *Schaalia odontolytica*, and demonstrate that each adhesin binds distinct components of the host cell wall. The adhesin from the motility driving T4P-1, termed PilX1, showed a wide range of affinities in microarray screening and bound peptidoglycan directly, but could not bind purified cell wall polysaccharides (CWPS) from host surfaces. Conversely, the adhesin from T4P-2, termed PilX2, bound host-derived CWPS and preferentially bound episymbiont-naïve hosts over episymbiont-exposed hosts. Within the bacterial host we identified two CWPS biosynthesis pathways that altered adhesin binding and episymbiont susceptibility.

Disruption of a rhamnose-CWPS pathway left hosts susceptible to episymbiont binding and disruption of a fucose remodeling pathway made hosts more resistant to episymbiont binding. Analyzing polysaccharide compositions revealed multiple CWPS, potentially including a rhamnose- and mannose-rich rhamnomannan. Consistent with this finding, exogenously added rhamnose and mannose were able to block binding by *N. lyticus*, supporting a role for these sugars in episymbiont defense. Our study reveals that Nanosynbacteria rely on the complex sugars that coat their host bacteria for recognition and attachment.

**Importance:** Minisyncoccota is a diverse phylum of ultrasmall bacteria that live in many environments, including the human microbiome, but remain understudied because they can only grow on the surface of specific prokaryotic hosts. This lifestyle is called episymbiosis and little is known about how episymbionts recognize and attach to their specific hosts. We found that *Nanosynbacter lyticus*, a model Minisyncoccota episymbiont, uses a pair of pilus associated proteins to mediate recognition and adhesion to molecules within hosts’ cell walls, including surface polysaccharides. Examining cell wall polysaccharide biosynthesis within the host revealed that changes in polysaccharide composition can increase or decrease susceptibility to episymbionts. These findings reveal distinct but overlapping adhesion roles for the two type 4 pili produced by *N. lyticus*, while also highlighting an important role for host surface sugars in influencing host-episymbiont compatibility.

## Introduction

All cultivated members of the phylum Minisyncoccota, also known as Patescibacteriota or the Candidate Phyla Radiation (CPR), are obligate episymbionts residing on other microbes (1–4). Despite being ubiquitous in various environments, including mammalian microbiomes, molecular mechanisms of bacterial host identification and binding amongst ultrasmall bacterial episymbionts remain largely unknown. Type 4 pili (T4P) are well conserved in this group and have been demonstrated to be vital for host association within the genetically tractable class Nanosynbacteria, also known as Saccharimonadia or TM7 (1, 5–7). T4P have been implicated in both host surface binding, and twitching motility of oral and insect associated Nanosynbacteria (6, 8). Members of the G1 clade of Nanosynbacteria (9), have evolved two distinct yet paralogous T4P filaments, T4P-1 and T4P-2 (6). As first-line host interaction mechanisms, these pili and their associated adhesins are important symbiosis factors and potential host range determinants. This study uses the model Nanosynbacterium *N. lyticus* (10) to investigate the binding specificity of T4P-dependent adhesins PilX1 and PilX2 on both hosts and host envelope components.

The cognate host of *N. lyticus*, *S. odontolytica*, has a specialized but largely uninvestigated cell wall that enables pleomorphic cell shapes, variable cell lengths, and multipolar branching. Previous research has demonstrated that host strains which have been experimentally evolved along episymbionts accumulate mutations in cell wall biosynthesis genes (11), linking envelope biosynthesis and episymbiosis. Furthermore, recent studies have identified a specialized episymbiont sensing and response (Esr) pathway within *Actinomycetes*, which defensively induces polysaccharide biosynthesis genes after episymbiont exposure (12).

*Actinomycetales* bacteria have been known to produce irregular cell wall polysaccharides (CWPS) containing 6-deoxyhexoses (6DH), such as rhamnose and fucose, for decades based on biochemical assays (13, 14), however these CWPS have rarely been investigated in detail.

Investigation of CWPS in agricultural pathogens from *Microbacteriaceae* indicates multiple polysaccharides within the peptidoglycan, analogous to teichoic acids (15, 16). For example, *Rathayibacter festucae* VKM Ac-1390T, produces three, a rhamnomannan (tetrasaccharide core Rha-Man-Rha-Man), a rhamnan (Rha repeat core), and a teichuronic acid (pentasaccharide core Man-Rha-Glc-Glc-Glc) (17). The number of polysaccharides and their structures vary between species, making them potential candidates for host-identification by episymbionts.

Using a combination of bioinformatic and genetic techniques we identified pathways for CWPS biosynthesis in *S. odontolytica*, partially analogous to rhamnan polysaccharide biosynthesis in *Streptococcus* species (18–20). We hypothesized that these polysaccharides would be receptor molecules for *N. lyticus* episymbiotic binding. We tested this by disrupting surface glycan biosynthesis within hosts, then assessing carbohydrate composition, susceptibility to *N. lyticus* infection, and susceptibility to T4P-dependent adhesin binding. Collectively our findings demonstrate that Nanosynbacteria utilize multiple adhesin pilins to move through the environment, identify hosts, and attach themselves to peptidoglycan and surface glycans.

Potential hosts in turn synthesize structural surface glycans that modulate episymbiont binding.

## Results

### T4P-1 and T4P-2 in *N. lyticus* both incorporate lectin-like minor pilins

Previous work identified two minor pilin genes within *N. lyticus* that encoded putative binding domains, *pilX1* and *pilX2*, which likely function as adhesins for their respective pili (6). We used AlphaFold3 (21) to predict each protein’s 3D structure, identified the C-terminal binding domains, and submitted them to DALI (22, 23) for structural similarity searching. The highest confidence PilX1 structural model (ipTM = 0.81) showed a jelly-roll domain with similarity to carbohydrate hydrolases, lectins, and a few bacteriophage proteins (Supplemental table 1). The best PilX2 structural model (ipTM = 0.92) was a heptabladed β-propeller similar to fucose binding lectins, carbohydrate hydrolases, DNA binding proteins, and peptidases (Figures 1A-B). Additionally, PilX2 had structural similarity to another known T4P adhesin protein, PilY1 from *P. aeruginosa*, which targets integrin glycoproteins (24).

**Figure 1.**
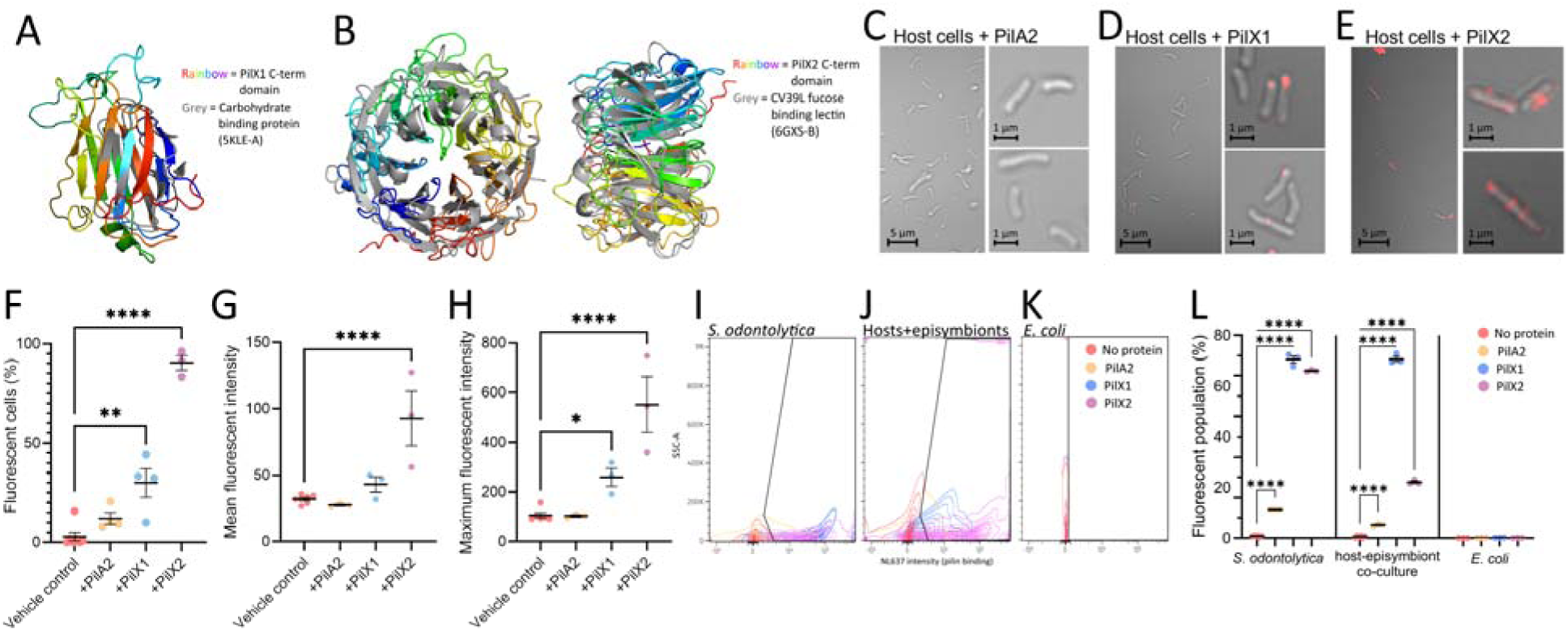
Episymbiont derived adhesins PilX1 and PilX2 bind host cell surfaces. **(A-B)** Structural similarity screening of T4P-dependent adhesin C-terminal domains via DALI suggested that **(A)** PilX1 had structural homology to carbohydrate binding/hydrolyzing proteins and **(B)** PilX2 was homologous to carbohydrate binding lectins and DNA binding proteins. Predicted AlphaFold3 structures (rainbow) are overlayed with the structures of their highest confidence DALI hits (grey). Representative phase contrast images of *S. odontolytica* cells labeled with **(C)** PilA2, **(D)** PilX1, or **(E)** PilX2. Labeled cells were quantified via **(F)** manually counting and FIJI quantification of **(G)** average and **(H)** maximum fluorescent intensity per cell. Flow cytometry of pilin-stained cells **(I-L)** allowed for direct comparison of pilin binding between **(I)** *S. odontolytica* monoculture, **(J)** host-episymbiont co-culture, and **(K)** the diderm control *E. coli*. Cytometry results are visualized as contour graphs with side scatter (SSC-A) on the y-axis and fluorophore intensity (NL637) on the x-axis. Unlabeled cells are shown in red, PilA2 in yellow, PilX1 in blue, and PilX2 in purple. Gates (bold black lines) were chosen so that ≈99% of untreated cells were negative for fluorescence. **(F-H, L)** Error bars indicate standard error of the mean. Significance determined via One-Way ANOVA with Dunnett’s test. * = p ≤ 0.05, ** = p ≤ 0.01, *** = p ≤ 0.001, **** = p ≤ 0.0001.

### PilX1 and PilX2 bind *S. odontolytica* host cells

To investigate the localization of PilX1 and PilX2 receptor molecules on host cells, putative binding domains were fused to a 6x His tag, a SUMO tag (for protein solubility), and a FLAG tag. The C-terminal domain from the major pilin PilA2 was expressed as a non-binding control. Fixed *S. odontolytica* cells treated with PilA2 showed little to no protein binding, similar to untreated cells (Figure 1C-H). *S. odontolytica* cells treated with PilX1 showed clear binding at distinct puncti, typically strongest at the cell poles (Figure 1D-H). *S. odontolytica* cells treated with PilX2 showed large amounts of bound protein at regularly spaced intervals across the cell (Figure 1E-H). Fluorescent cells were counted manually to determine binding frequency (Figure 1F). Since both proteins showed punctiform binding rather than diffuse binding, image analysis was used to calculate mean (Figure 1G) and maximum (Figure 1H) signal intensity. Analysis supported moderate host binding by PilX1 and strong host binding by PilX2.

To quantitatively describe binding, flow cytometry was performed to quantify adhesin bound host populations (Supplemental table 2). Using uninfected hosts, we found that ≈11.3% of cells treated with PilA2, ≈70.5% of cells treated with PilX1, and ≈65.9% of cells treated with PilX2 were more fluorescent than untreated cells (Figures 1I, 1L). While no PilA2 binding had been detected via microscopy, flow cytometry suggested that PilA2 may sub-visually bind *S. odontolytica*. Since infection by episymbionts is known to significantly alter host morphology and metabolism, adhesin binding was repeated with a stable co-culture of host cells and episymbionts. Examining particles within the host cell size range revealed similar binding frequency by PilA2 (≈5.2%) and PilX1 (≈70.6%); however, PilX2 bound co-culture cells significantly less (≈22.1%) than uninfected hosts (Figures 1J, 1L). Reduced PilX2 binding in co-cultures could indicate changes to the host cellular envelope after infection or steric blocking of the adhesin’s receptor molecule by episymbionts. Within co-cultures, PilX1 also induced aggregation, resulting in decreased particle counts and a high side scatter population (>1 million a.u.) that was absent in in other treatments (Figure 1J), indicating that PilX1 may crosslink cells *in vitro*. No protein effectively bound the diderm bacterium *E. coli* (Figures 1K, 1L).

### Glycan and peptidoglycan screens reveal distinct binding affinities for PilX1 and PilX2

To experimentally screen PilX1 and PilX2 for binding epitopes, purified protein was applied to a microbial glycan microarray (MGM) containing 313 unique microbially-derived polysaccharides (Figures 2A-C, Supplemental table 3). PilX1 displayed low to moderate binding on a wide range of glycans, with the strongest signals coming from O-antigen polysaccharides (OPS) from *Proteus mirabilis* strains O27 (25) and O41 (26), *Shigella flexneri* type 1a (27), and *Shigella boydii* type 13 (28) (Figure 2B). Comparing the structures of these glycans (Figure 2D) reveals that the only unifying feature is the presence of N-acetyl glucosamine (NAG). The glycan with the strongest affinity specifically had NAG within its linear backbone and as a decoration of that backbone. The breadth of PilX1 binding partners suggests that this adhesin may bind multiple receptors or binds a commonly occurring epitope. PilX2 showed little to no binding on any of the assayed glycans (Figures 2A, 2C). This could indicate either that PilX2 does not bind a glycan moiety, requires a cofactor to initiate binding, or only binds an epitope that is not present on the microarray. Microarray analysis is useful for high throughput screening, however the range of microbially-derived glycans is limited and heavily biased towards Gram-negative human pathogens (290 out of 313). The MGM only contains 23 glycans from Gram-positive bacteria, and they are all capsular polysaccharides from *Streptococcus pneumoniae* rather than CWPS. Next, all three proteins were applied to pure peptidoglycan from *B. subtilis* and isolated cell walls from host *S. odontolytica* via immune-dot blotting (Supplemental figure 1). Both PilX1 and PilX2 strongly bound the pure peptidoglycan (Figure 2E), however the host-derived cell wall sacculi showed reduced PilX2 binding, and no significant PilX1 binding (Figure 2F). This suggests that both proteins can bind peptidoglycan, however some component of the host cell walls, potentially CWPS, reduces access to peptidoglycan.

**Figure 2.**
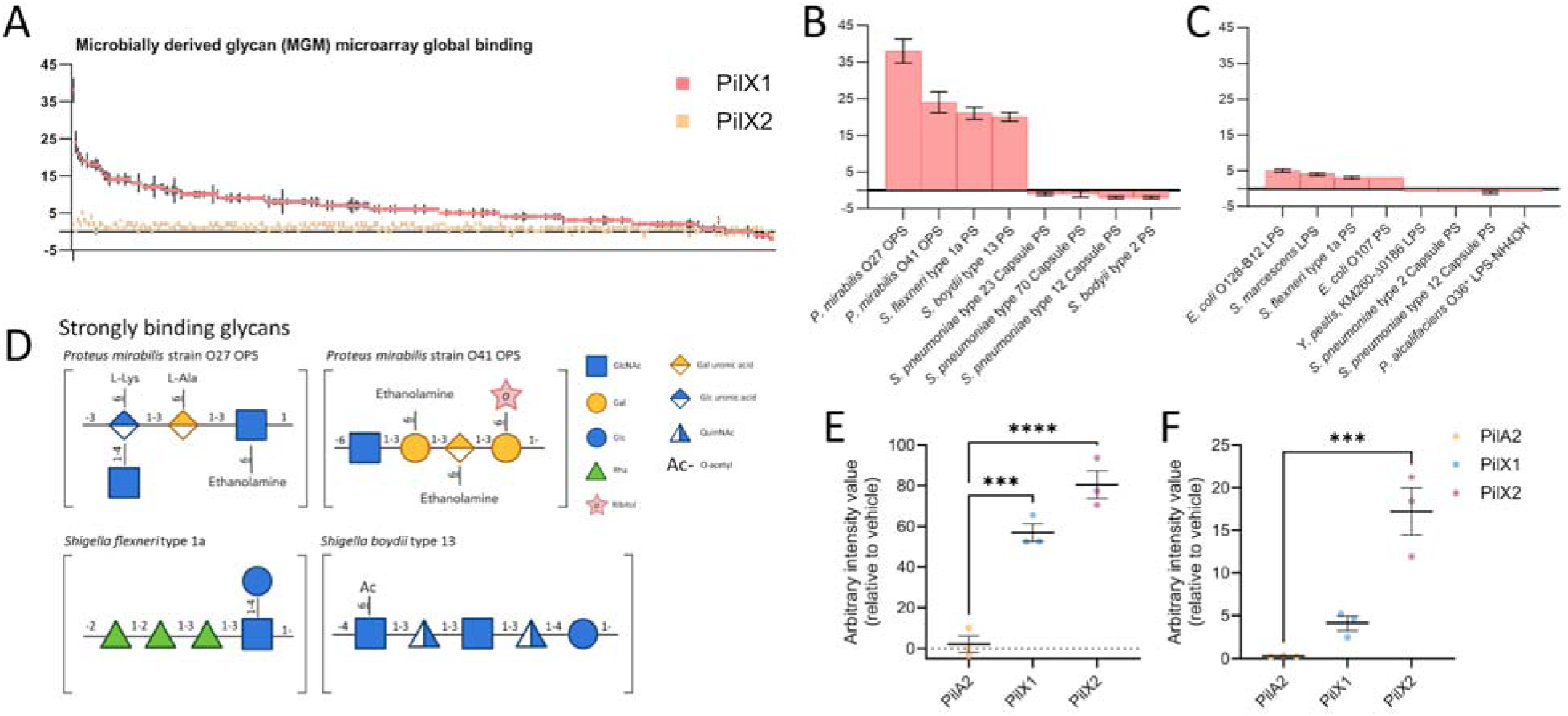
T4P-dependent adhesins have distinct binding affinities for glycans and peptidoglycan. **(A-C)** Applying our two adhesin proteins to microarrays containing 313 purified glycans revealed distinct binding behavior. Individual glycans are shown on the x-axis, relative fluorescence units are on the y-axis, and error bars depict the standard error of the mean. Graph **(B)** shows the four glycans with the strongest and weakest PilX1 binding signals, graph **(C)** shows the four strongest and weakest PilX2 binding glycans. **(D)** Examining the PilX1 binding glycans reveals diverse structures, however they all incorporate N-acetyl glucosamine. **(E)** Immuno-dot blotting pure peptidoglycan showed strong binding by both adhesins, however **(F)** whole cell walls extracted from the host organism, *S. odontolytica*, only showed significant binding by PilX2. **(E-F)** Error bars show standard error of the mean. Significance determined via One-Way ANOVA with Dunnett’s test. * = p ≤ 0.05, ** = p ≤ 0.01, *** = p ≤ 0.001, **** = p ≤ 0.0001.

### Disruption of host CWPS biosynthesis alters episymbiont susceptibility

To elucidate if and how host-derived CWPS alter episymbiont susceptibility, knockout mutations were attempted in two distinct CWPS biosynthesis pathways. First, we targeted a *marR*-like regulatory protein (APY09_RS07555, henceforth *marR*) and a fucosidase/fucose uptake operon (APY09_RS05345 carbohydrate binding protein; APY09_RS05350 ABC transporter permease; APY09_RS05355; fucosidase) predicted to coordinately play roles in glycan remodeling (Supplemental figure 2A). Consulting a transcriptomic dataset of episymbiont exposure (7) shows that this regulatory protein and the genes within the FucoseLocus are inversely affected during host exposure, with downregulation of the predicted regulator, and upregulation of the predicted fucose transporter genes (Figure 3A).

**Figure 3.**
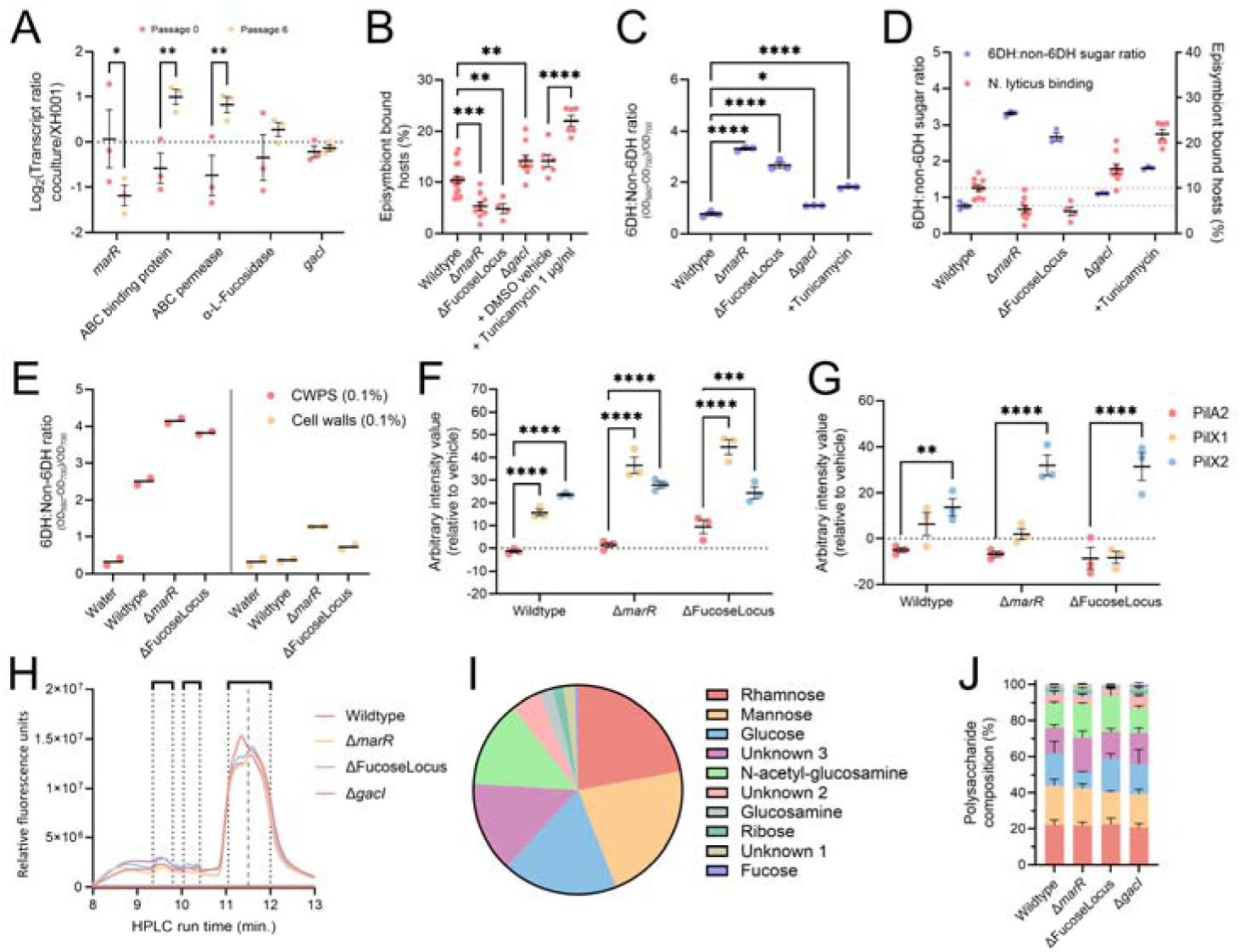
*S. odontolytica* produces complex surface glycans, and disruption of glycan biosynthesis modulates episymbiont binding. *S. odontolytica* encodes many genes associated with polysaccharide assembly, decoration, and recycling. **(A)** Examining a previously published transcriptomic dataset (7), reveals a MarR-family regulator and a fucose recycling locus that are differentially expressed after episymbiont exposure (co-culture passage 6) relative to pre-exposure (co-culture passage 0). Expression of the transglycosylase *gacI* was not episymbiont responsive. **(B)** CWPS biosynthesis mutants, as well as cells treated with the CWPS chemical inhibitor tunicamycin, were infected with *N. lyticus*, revealing both increased and decreased susceptibility. **(C)** A modified anthrone assay was used to calculate the ratio of 6DH (rhamnose and fucose) to non-6DH sugars, and all mutants had elevated 6DH compositions **(D)** despite having inverse binding phenotypes. To characterize the host CWPS, glycans were cleaved from cell walls and purified. An anthrone assay **(E)** supported elevated 6DH content within Δ*marR* and ΔFucoseLocus mutants’ CWPS. Dot blots were prepared with **(F)** glycan depleted cell walls and **(G)** the isolated CWPS fractions, and then immunostained with labeled PilA2 (red), PilX1 (yellow), or PilX2 (blue). **(H)** Isolated CWPS were separated via HPLC to examine polysaccharide content. Size exclusion chromatography showed the presence of ≈5 polysaccharides, with run time of elution shown on the x-axis and fluorescence on the y-axis. Next, polysaccharide fractions were hydrolyzed and separated via HPAEC-PAD; Graph **(I)** is a pie diagram visualizing the ratio of monosaccharides present in wildtype and graph **(J)** compares the composition of CWPS biosynthesis mutants side by side. **(A-D, F-G, J)** Error bars show standard error of the mean. Significance determined via One-Way ANOVA with Dunnett’s test. * = p ≤ 0.05, ** = p ≤ 0.01, *** = p ≤ 0.001, **** = p ≤ 0.0001.

Secondly, genomic investigation of *S. odontolytica* identified many homologs of rhamnose-rich CWPS (Rha-CWPS) biosynthesis proteins originally characterized in *Streptococcus* systems. Within Streptococcal species, Rha-CWPS act like wall teichoic acids (29), contribute to phage and host interactions (19), and can be used to identify immunological serotypes (20). Within *S. odontolytica*, these genes are found in five genomic loci and include dDTP-rhamnose synthesis (*rml* genes), a rhamnose-fucose epimerase, Rha-CWPS extension and export (*rgp* genes), up to seven predicted glycosyltransferases, and a system predicted to add phosphocholine decoration to teichoic acids, based on analogy to *S. pneumoniae* (30, 31) (Supplemental figure 2B).

After thorough analysis, key genes were picked and homologous recombination constructs were used to target *marR*, the FucoseLocus, *rmlC-D* (APY09_RS00100), *rgpB* (APY09_RS05560), *rgpF* (APY09_RS05510), *tagO* (APY09_RS06605), a putative glycosyltransferase (APY09_RS05455), a putative N-acetyl glucosyltransferase (APY09_RS00090), and a convergently encoded neutral site (between APY09_RS05760 and APY09_RS05765). Only *marR*, the FucoseLocus, the putative N-acetyl glucosyltransferase (annotated as *gacI*), and the neutral site could be successfully mutated, indicating that almost all the Rha-CWPS associated genes were essential for survival, consistent with a previously performed transposon library screen (32). To overcome this limitation, a chemical inhibitor of TagO, tunicamycin, was also used to disrupt Rha-CWPS biosynthesis (18, 20). Growth curves were performed to determine an optimal concentration. Microscopic examination of tunicamycin treated co-cultures showed abundant episymbiont growth at all tested concentrations, even those with strongly inhibited host growth, suggesting that tunicamycin is less cytotoxic to the episymbiont than the host (Supplemental figure 3A). A concentration of 1.2 μM (1 μg/mL) was chosen to allow *S. odontolytica* growth while maximizing cell wall impact (Supplemental figures 3B-C).

All CWPS biosynthesis mutants, and tunicamycin treated cells, were infected with episymbionts to assess binding potential (Figure 3B). ≈10.4% of wildtype host cells were bound within the first 24 h. Binding by episymbionts was reduced on Δ*marR* (≈5.4%) and ΔFucoseLocus cells (≈4.9%); however, binding was enhanced on Δ*gacI* (≈14.2%) cells.

Tunicamycin treated cells (≈22.0%) had increased binding relative to vehicle control (≈14.2%). Deletion of *marR* or the FucoseLocus also resulted in significantly shorter cell sizes, connecting glycan remodeling to cell morphology (Supplemental figure 4A). A modified anthrone assay was used to understand how CWPS compositions differed between genotypes (33). This assay converts sugars into chromophores to provide a relative abundance ratio of 6-deoxyhexoses (6DH) relative to non-6DH sugars, while having no effect on NAG/NAM residues from peptidoglycan (Supplemental figure 4B). Since fucose and rhamnose are both 6DH, and most other polysaccharide constituents are not, the anthrone assay should theoretically detect compositional changes from disruption of either CWPS pathway. Cell walls from wildtype *S. odontolytica* had a 6DH ratio of ≈0.77, while the Δ*marR* mutant (≈3.30), ΔFucoseLocus mutant (≈2.67), Δ*gacI* mutant (≈1.10), and tunicamycin treated cells (≈1.81) had significantly higher 6DH ratios (Figure 3C). These results indicate that disruption of the *marR*-associated fucosidase pathway results in large 6DH increases that block episymbiont binding, while disruption of the Rha-CWPS pathway leads to minor 6DH increases that increase hosts susceptibility to episymbiont binding (Figure 3D).

### Isolation and characterization of *S. odontolytica* CWPS

Since disruption of both CWPS biosynthesis pathways altered episymbiont binding and CWPS composition (Figures 3B-3D), pure polysaccharide fractions were prepared for analysis, focusing on the most strongly impacted strains, Δ*marR* and ΔFucoseLocus. Trichloroacetic acid cleavage was used to remove CWPS from peptidoglycan (and other insoluble components) (34). CWPS and CWPS-depleted cell wall fractions were then analyzed via anthrone assay to determine TCA cleavage efficacy. For wildtype *S. odontolytica*, 75.9% of reactive saccharides were in the CWPS fraction (Supplemental figure 4C). For *S. odontolytica* Δ*marR* and ΔFucoseLocus 84.8-86.3% of the reactive saccharides were in the CWPS fraction, suggesting that mutants were either less resistant to glycan hydrolysis or had fewer glycans to be cleaved.

Purified mutant CWPS still showed elevated 6DH, reflecting the whole cell wall anthrone assay (Figure 3E). However, the wildtype CWPS fraction had a greater 6DH ratio than whole cell walls, suggesting that the CWPS that remained bound to peptidoglycan after hydrolysis may disproportionately contain non-6DH sugars.

To assess PilX1 and PilX2 binding of pure CWPS and CWPS-depleted peptidoglycan, 0.1% (w/v) solutions of all fractions were spotted onto nitrocellulose membranes to assess pilin adhesin binding via dot blot immunostaining (Supplemental figure 4D). PilA2 was used as a binding control, showed little binding on depleted cell wall fractions, and was actively repelled by CWPS fractions (Figures 3F-G). Despite PilX1 not significantly binding whole cell walls (Figure 2F), PilX1 did significantly bind all CWPS-depleted cell wall fractions. Notably, it bound mutant cell walls more strongly than wildtype. As our prior anthrone assay indicated, depleted wildtype cell walls retained more glycans than mutants cell walls (Supplemental figure 4C), so this pattern suggests that uncleaved wildtype glycans may still partially inhibit PilX1 binding. PilX1 did not bind CWPS fractions from any *S. odontolytica* genotypes (Figures 3F-G).

PilX2 shows a quite different binding profile. PilX2 bound all CWPS-depleted cell wall fractions similarly, supporting its potential for peptidoglycan binding. PilX2 also moderately bound wildtype CWPS and strongly bound *S. odontolytica* Δ*marR2* and ΔFucoseLocus CWPS fractions (Figures 3F-G). This increased PilX2 binding was unexpected, since these mutants have decreased susceptibility to episymbiont binding. This inverse binding could indicate that the decrease in episymbiont susceptibility results from blocking of PilX1 or a yet undiscovered adhesin rather than blocking of PilX2. Additionally, the Δ*marR2* and ΔFucoseLocus mutants have significantly shorter cell lengths, so host surface area may also affect susceptibility.

Collectively, these data paint a picture of a complicated assembly of surface glycans that can block PilX1 adhesion, be bound directly by PilX2, and holistically modulate susceptibility to episymbiont infection. Despite both associating with pure peptidoglycan, PilX1 and PilX2 appear to have distinct receptors indicative of distinct, if overlapping, binding functions.

To better understand CWPS content, purified CWPS fractions were analyzed via HPLC to quantify constituent glycans. Using size exclusion chromatography to separate polysaccharides, we identified five peaks/polysaccharides present in all assayed samples (Figure 3H). A small peak centered around 9:30 min, two small peaks (combined for analysis) around 10:15 min, and a broad peak around 11:30 comprising two partially resolved components (separated for analysis) (Supplemental figure 4E). The components producing the three early eluting peaks were present at lower abundance (∼10.9-17.5%) than the components producing the broad, partially resolved peak (∼82.5-89.1%). Traces were similar between genotypes; however, for wildtype CWPS the leading component of the broad peak was predominant, whereas in all mutants the trailing component was predominant.

Purified CWPS fractions were then hydrolyzed into constituent monosaccharides and separated via HPAEC-PAD to determine compositions. HPAED-PAD detected ten components in wildtype *S. odontolytica* CWPS, seven of which could be identified via standards (Figure 3I, Supplemental figure 4F). The most prevalent components were rhamnose (22.1%) and mannose (21.9%), being roughly equimolar. This may suggest that they occur as a rhamnomannan rather than a classical Rha-CWPS. Glucose (17.9%), unknown 3 (14.0%), and NAG (13.4%) were also abundant. The remaining components, including fucose (0.4%), all had less than 5% abundance. Comparing monosaccharide ratios between genotypes revealed surprisingly similar monosaccharide profiles, though Δ*marR* had significantly decreased glucose, and Δ*marR* and ΔFucoseLocus trended towards increased NAG (Figure 3J, Supplemental figure 4G), potentially explaining their altered PilX1 and PilX2 binding profiles (Figures 3F-3G).

### Exogenous sugars modulate episymbiont binding

Since CWPS compositional analysis could not identify which component(s) inhibited episymbiont binding, *N. lyticus* was added to cultures of *S. odontolytica* supplemented with exogenous monosaccharides present within CWPS (15–17). Mannose had to be applied at a lower concentration because at high concentrations mannose inhibited host growth (Supplemental figure 5), suggesting sugar toxicity or accumulation of toxic intermediates (35). *N. lyticus* binding in the absence of sugars (14.3 ± 1.5%) was not significantly different from sucrose (13.3 ± 1.3%) or fucose (12.8 ± 3.3%). However, binding was significantly decreased by rhamnose (1.2 ± 0.8%), glucose (6.3 ± 1.5%), and low-concentration mannose (7.7 ± 1.5%) (Figure 4A). A T4P chemical inhibitor known to reduce binding, quercetin, significantly decreased binding (4.3 ± 1.1%), but not as strongly as rhamnose. A rhamnose concentration response curve was generated between 5 mM and 200 mM, measuring frequency of host-binding (Figures 4B-C) and total *N. lyticus* (Figure 4D). Concentration dependent inhibition of host-binding was observed with an inhibition coefficient (IC50) of 19.77 mM, despite there being no significant decrease in total episymbiont population. To determine if rhamnose-dependent binding inhibition was driven by T4P-2 dependent PilX2 binding, three species of *Nanosynbacteraceae* that encode similar PilX2 homologs, *N. lyticus*, HMT-488 strain BB004 (PilX2 with 77% aa identity), and HMT-352 stain TM7-008 (PilX2 with 69% aa identity) were grown in exogenous rhamnose (6, 36, 37). TM7-008 displayed decreased binding comparable to *N. lyticus*, while there was no significant impact on BB004 binding (Figure 4E). These data suggest that the rhamnose and mannose present in *S. odontolytica*’s CWPS are protective against some episymbiont species, but T4P-2 and its adhesin PilX2 are unlikely to be the mechanism of inhibition.

**Figure 4.**
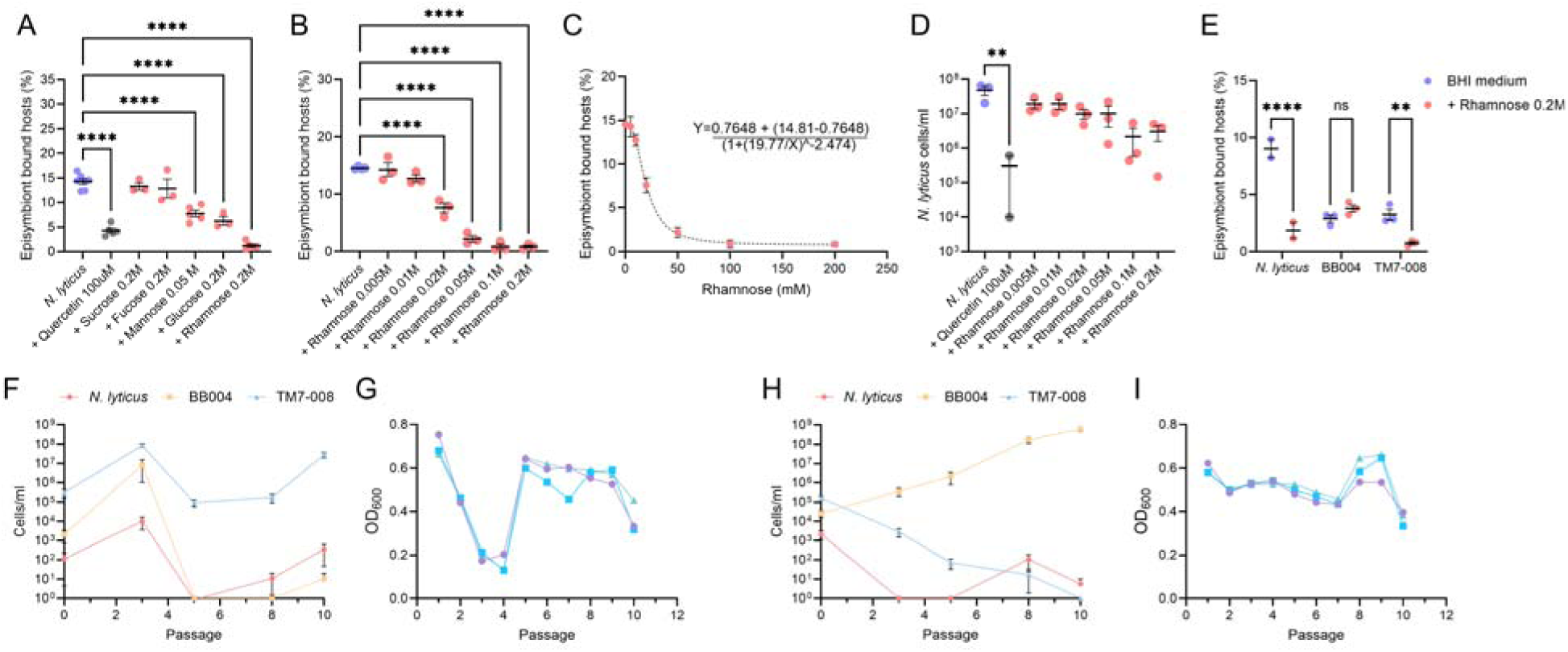
Exogenous carbohydrates inhibit episymbiont binding, in a species dependent manner. **(A)** Host episymbiont co-cultures established in various monosaccharides identified three sugars capable of inhibiting binding. **(B)** Rhamnose had the greatest impact, displaying exponential dose dependent inhibition **(C)** with an IC50 of ∼20 mM. **(D)** Despite ablation of host binding, total TM7x counts were not significantly decreased. **(E)** Examining three Nanosynbacteria capable of infecting the same host species revealed that two were affected by rhamnose-dependent inhibition, TM7x and TM7-008, while the third was unaffected. A mock community containing all three Nanosynbacteria was allowed to grow across ten serial passages, in the absence **(F-G)** or presence **(H-I)** of exogenous rhamnose. Graphs **(F)** and **(H)** show relative abundances of each episymbiont as determined via qPCR. Graphs **(G)** and **(I)** show the optical densities of mock communities, with each biological replicate shown in a different color. **(A-I)** Error bars show the standard error of the mean. Where shown, significance determined via One-Way ANOVA with Dunnett’s test. * = p ≤ 0.05, ** = p ≤ 0.01, *** = p ≤ 0.001, **** = p ≤ 0.0001.

### Nanosynbacteria species competition is modulated by sugars

Previous work has demonstrated that binding defects, like those induced by the loss of T4P-2, alter the competitive fitness of an episymbiont like *N. lyticus* (6, 38). To determine if rhamnose-dependent binding inhibition also impacted competitive fitness, we utilized tripartite competition between the above three *Nanosynbacteraceae* episymbionts. When passaged in un-supplemented medium, host-populations collapsed during passage 3 and the previously reported hierarchy of fitness was observed (6), TM7-008 > *N. lyticus* > BB004 (Figures 4F-4G).

Conversely, when these species were serially passaged in medium supplemented with 50 mM L-rhamnose, the host crash never occurred, and the competitive hierarchy was inverted, BB004 > *N. lyticus* > TM7-008 (Figures 4H-4I). These data suggest that rhamnose-dependent inhibition of episymbiont binding is species specific, highlighting how sugar-dependent inhibition, and potentially CWPS-dependent inhibition, can have drastic implications for population dynamics.

### T4P-1 drives twitching motility, which is induced by host proximity

Despite CWPS blocking PilX1 binding, this protein was still found to associate with peptidoglycan, host cells, and a range of glycan microarray candidates; suggesting PilX1 may have a broad range of receptor molecules. Previous studies suggested that T4P-1, associated with PilX1, was responsible for twitching motility, a function which may benefit from being able to bind diverse surfaces (6). However, this role could not be tested previously because T4P-1 genes are essential for survival. We generated a T4P-1 deficient strain by knocking down expression of its major pilin component, PilA1, via genomic integration of an antisense *pilA1* transcript cassette (*N. lyticus* NS1::Antisense-*pilA1*). Knockdown efficacy was determined via rt-qPCR, revealing a 91.8% reduction in expression of *pilA1* (Figure 5A). Using timelapse phase contrast microscopy, the motion of planktonic episymbionts within co-cultures was tracked (Figure 5B). Our control, a neutral site insertion mutant, had an average speed of 124.0 nm/sec. and a T4P-2 defective strain (*N. lyticus* Δ*pilA2*) showed no decrease in twitching motility (∼141.0 nm/sec.), consistent with previously results (6). Conversely, the T4P-1 defective strain showed almost total motility inhibition (∼22.3 nm/sec.), only slightly more mobile than heat-killed cells (15.1 nm/sec.) (Supplemental video 1).

**Figure 5.**
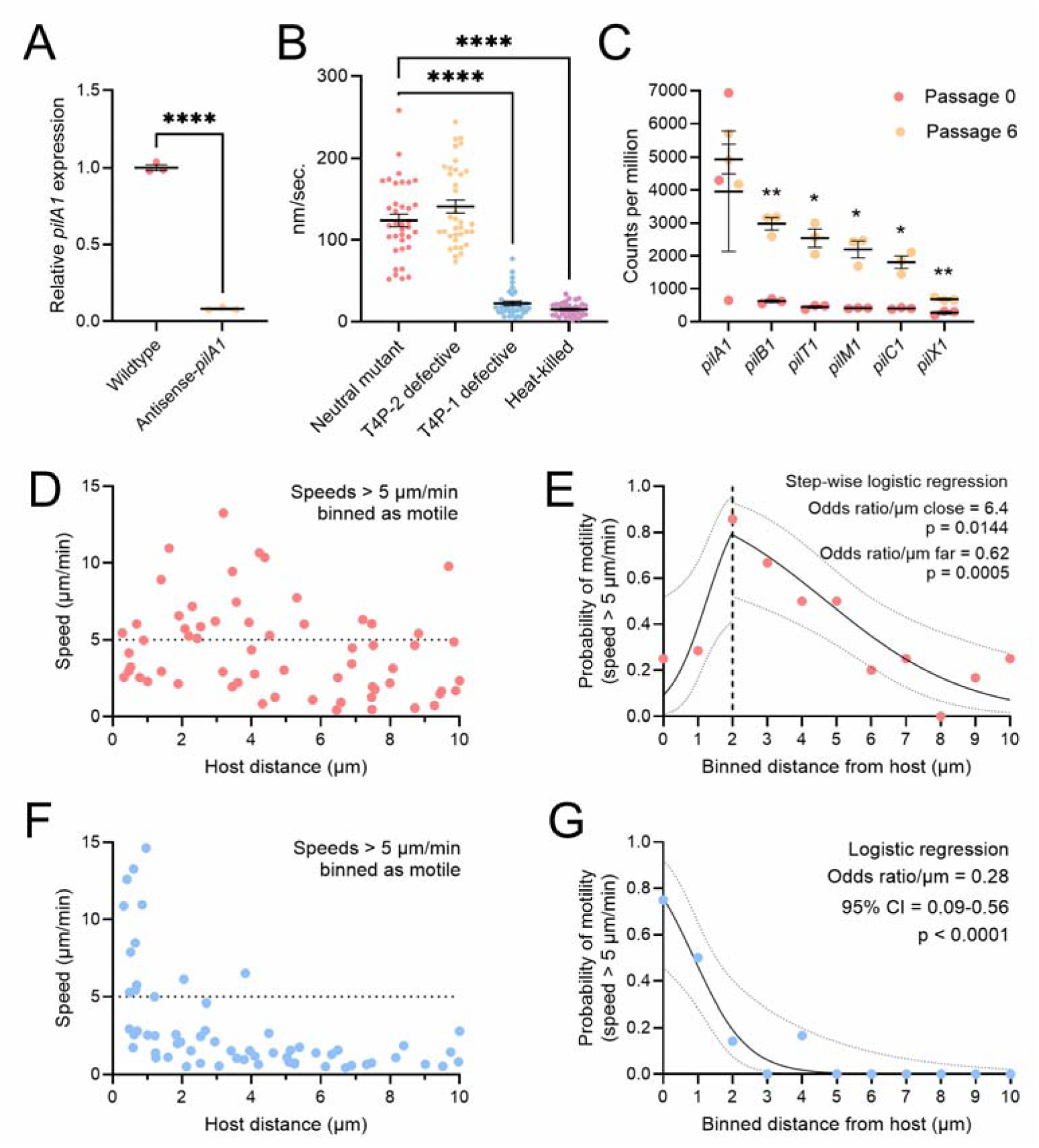
Twitching motility is T4P-1 dependent and induced by host proximity. During its planktonic lifecycle, *N. lyticus* cells utilize T4P-dependent twitching. **(A)** To determine if T4P-1 is required for twitching motility, a knockdown mutant for its major pilin, *pilA1*, was generated, resulting in a 91.8% transcriptional reduction. **(B)** Time lapse imaging (Supplemental video 1) was used to calculate average twitching speeds. Wildtype cells moved at ≈120 nm/sec. Deletion of the T4P-2 major pilin, PilA2, had no effect on twitching speed. Knockdown of the T4P-1 major pilin, PilA1, drastically reduced motility, almost to the level of heat-killed cells. **(C)** Previously conducted transcriptomics (7) show that most T4P-1 genes within *N. lyticus* are transcriptionally upregulated after host infection (passage 6) relative to pre-infection (passage 0). Timelapse microscopy was used to calculate twitching speed as a function of host-distance for both wildtype **(D-E)** and *pilA1* knockdown cells **(F-G)** to look for proximity induction. Graphs **(D)** and **(F)** show the speeds of individual cells (y-axis) as a function of their host-distance (x-axis). Graphs **(E)** and **(G)** bin cells as motile (>5 μm/min) or non-motile to describe the probability of motility via logistic regression. **(A-C)** Significance determined via One-Way ANOVA with Dunnett’s test. * = p ≤ 0.05, ** = p ≤ 0.01, *** = p ≤ 0.001, **** = p ≤ 0.0001.

Visually, T4P-1 defective cells only twitched when located close to host bacteria, potentially suggesting that proximity induced enough *pilA1* expression to overcome knockdown. Supporting this idea, a previously published transcriptomic dataset (7) shows that most T4P-1 genes are upregulated after host exposure (Figure 5C). Quantifying cell speeds (N=63) as a function of initial host-distance revealed that wildtype episymbiont motility increased with proximity, but only up to a point. Wildtype *N. lyticus* were most motile ≈2 μm from their host (Figures 5D-E), cells closer than that showed reduced movements, potentially due to initiation of host-surface attachment. Strikingly, T4P-1 defective *N. lyticus* showed wildtype levels of motility between 0-1 μm host-distance. Beyond this distance, motility rapidly declines, with no visible motility past 4 μm host-distance (Figures 5F-G). These data show that episymbionts increase speed when near hosts and suggest that T4P-1 expression contributes to proximity induced motility.

## Discussion

Characterization of PilX1 from T4P-1 and PilX2 from T4P-2 revealed clear cell wall association on host cell surfaces, supporting host binding by both T4P-1 and T4P-2. Since these proteins resembled lectins, we investigated the poorly characterized surface polysaccharides produced by the episymbiont’s host as potential receptor molecules (39, 40). Combining glycan microarray, peptidoglycan, and CWPS binding results showed that both proteins bind peptidoglycan, but host cell wall binding, particularly by PilX1 binding, is reduced by the presence of cell wall modifications. To determine if CWPS decoration is responsible for adhesin blocking, we genetically targeted pathways for fucose remodeling/uptake and rhamnose-rich polysaccharide synthesis, resulting in mutants with distinct CWPS compositions and susceptibility to episymbiont binding. These results demonstrate that CWPS are potentially defensive structures within host cell walls, whose architecture can dictate episymbiont binding. This is supported by a recent study which demonstrates that *Actinomycetes* encode Esr signaling systems which detect episymbionts and defensively induce expression of polysaccharide biosynthesis pathways, as well as secreted defense factors (12). However, disruption of CWPS pathways also altered cell sizes, suggesting that polysaccharides also play structural roles.

PilX1 showed no binding on any CWPS fractions but was able to bind cell walls after CWPS was removed, suggesting that PilX1’s primary receptor is peptidoglycan. Conversely, PilX2 bound some component(s) of the CWPS, and binding was enhanced in fucose remodeling/uptake mutants despite those mutants having reduced susceptibility to episymbiont binding. Based on these data we hypothesize that PilX2 binds an epitope within the CWPS, potentially a fucose residue since deletion of the fucosidase locus increased binding. Despite increased adhesin binding in these mutants, episymbiont susceptibility decreased. Perhaps this mutant is less susceptible to episymbiosis as a result of i) blocking a yet undiscovered adhesin, ii) decreased surface area from shorter cell lengths, or iii) broad disruption of cell envelope architecture.

Examining the composition of *S. odontolytica*’s CWPS revealed ∼5 polysaccharides with abundant and equimolar rhamnose and mannose, suggesting that one of these polysaccharides may be a rhamnomannan (Figure 5I), like the rhamnomannan produced by some *Microbacteriaceae* (16). The three most common CWPS components, rhamnose, mannose, and glucose, were the same monosaccharides that inhibited *N. lyticus* binding when added to cultures exogenously, supporting the idea that these sugars act to mask receptor sites or otherwise suppress episymbionts. Despite mutation of a fucose remodeling locus impacting episymbiont binding, fucose itself was the least abundant monosaccharide detected. This suggests that fucose may be a rare or apical decoration of host polysaccharides.

PilX1 and PilX2’s distinct binding behaviors are particularly interesting in the context of their associated pili. Our experiments demonstrated that T4P-1 is essential for twitching motility in *N. lyticus*. Serendipitously, we found that host proximity induces increased episymbiont motility. Since proximity induced motility was most significant in *pilA1* knockdown cells, upregulation of T4P-1 genes may prominently contribute to proximity induced motility.

Additionally, since T4P-1 associated PilX1 binds peptidoglycan and a range of glycan microarray constituents, it appears to be a relatively generalist binding machine. This could be beneficial for a motility adhesin that needs to be able to interact with diverse organic surfaces to navigate biofilms and epithelial surfaces. Conversely, T4P-2 is known to contribute to host binding and does not play a role in twitching motility (6, 38). In line with a host binding function for T4P-2, its adhesin, PilX2, bound a yet unknown CWPS present on host surfaces. Interestingly, PilX2 bound episymbiont-naïve host cells approximately three times more than it bound episymbiont infected host cells, suggesting that T4P-2 allows Nanosynbacteria to avoid binding episymbiont burdened hosts. This observation could suggest; i) superinfection may be unfavorable because an episymbiont burdened host provides fewer nutrients, ii) host bacteria could modify CWPS in response to episymbiont exposure (12), or iii) prior infection by *N.lyticus* removes or sterically blocks host receptor sites. Superinfection inhibition mechanisms have been previously observed in bacteriophage (41, 42), where they are hypothesized to maintain parasite diversity.

One limitation of this study is that we were unable to determine definitive binding epitopes for PilX1 and PilX2. Future studies should i) use characterized peptidoglycan fragments to identify which are bound by each adhesin and ii) purify individual polysaccharides from CWPS fractions to identify which associates with PilX2. Synthesizing these data into a potential model, we predict that *S. odontolytica* produces multiple CWPS, which contribute to cell shape and mask peptidoglycan, reducing binding by generalist adhesins like PilX1; however, *N. lyticus* has adapted to bind one of these CWPS using PilX2 (24, 43) (Figure 1B). PilX2 may also enable *N. lyticus* to preferentially bind uninfected hosts as a superinfection avoidance strategy. Regulatory proteins like the identified *marR*-family regulator enable hosts to modulate their surface glycans in response to episymbiont exposure. Considering the ancient co-evolutionary Nanosynbacteria-Actinobacteria relationship, species-specific surface glycans have likely been shaped by predator-prey dynamics. Within our model, Actinobacteria metaphorically act as arborists, carefully populating a forest of glycans on their surface to balance structural needs with ectoparasite resistance, meanwhile Nanosynbacteria have adapted adhesins and binding appendages to find compatible glycan groves that they can detect and interact with.

## Materials and Methods

### Bacteriological culture conditions

Bacterial strain list present in Supplemental table 4. Cloning and expression strains of *E. coli* were grown in LB medium at 37°C. All other broth cultures were grown in reduced brain-heart infusion medium (BHI) in a microaerophilic Whitley A35 workstation at 37°C. Microaerophilic cultures were grown in 2% O_2_, 5% CO_2_, balanced with N_2_. Nanosynbacteria were stored at -80°C in 15% Glycerol to maintain viability. To standardize the multiplicity of infection (MOI) for competitions, qPCR primers #1-6 (Supplemental table 5) were used to quantify Nanosynbacteria species (6). For episymbiont binding experiments, a previously published OD_600_ standard curve was used to quantify Nanosynbacteria (6). Growth curves were performed using a Cerillo (Charlottesville, VA) Stratus microplate reader.

### Bioinformatic analyses

Where identified, genomic loci were visualized in Geneious Prime 2024.0.7. SignalP v6.0 (44) was used to predict signal peptides. To gauge expression of the genes within the MarR regulome, transcriptomic reads/million data was extracted from a previously published dataset (GEO GSE196744) (7). For structural alignments, the highest confidence AlphaFold3 (45) models for PilX1 and PilX2 were trimmed to their binding domains in PyMol 3.0.0 (46). Resulting structures were analyzed via DALI using the PDB25 search with default parameters (22). Outputs with roles in carbohydrate/glycan hydrolysis or binding were highlighted (Supplemental table 1). Carbohydrate binding proteins of from each DALI output (5KLE and 6GXS) were aligned against the query structure using the RCSB protein Data Bank alignment tool and the jFATCAT (rigid) method (47–50).

### Design and expression of pilin proteins

As informed by the Conserved Domain Database (51), pilin encoding sequences were trimmed to just include predicted binding domains (PilA2 = Y34-Ochre142; PilX1 = R32-Ochre381; PilX2 = P159-Ochre577), cloned into pET22b(+), and N-terminally fused a 6xHis tag for purification, a SUMO tag for enhanced solubility, and a FLAG tag using primers #7-25.

Plasmids were transformed into Bl21DE3. Cells were grown into midlog phase in BHI medium supplemented with 100 μg/ml ampicillin, chilled to 16°C, and induced with 0.25 mM Isopropyl β-D-1-thiogalactopyranoside (IPTG) for ≈16 h. To enhance the solubility of the relatively insoluble PilX1 and PilX2 constructs, a urea-based strategy was used (52). Briefly, harvested cells were re-suspended in solubility buffer (2 M urea, 300 mM NaCl, 100 mM tris-HCl, pH 12), sonicated at 30% amplitude (375 Vrms) for 20 min. and the resulting supernatant was mixed 50/50 with gentle renaturing buffer (1 M urea, 150 mM NaCl, 50 mM tris-HCl, 5% glycerol, 5% sucrose, pH 8). Proteins were purified via TALON SuperFlow resin (1-2 ml), eluted with 200 mM imidazole, dialyzed with wash buffer (300 mM NaCl, 50 mM NaH_2_PO_4_, pH 7) twice, and dialyzed with storage buffer once (50/50 wash buffer/glycerol). Protein concentration was determined via Peirce 660 nm reagent (1861426) or via Micro BCA kit (ThermoScientific 23235) prior to storage at -80°C.

### Pilin labeling

Samples were grown to late log phase in triplicate, chilled to arrest growth, fixed in a 5% formaldehyde PBS solution for 15 min. at ambient temperature, and subjected to a light lysozyme treatment (5 min. at 2 μg/ml) to expose cell wall epitopes. Fixed cells were labeled with 50 μg of protein (PilA2, PilX1, or PilX2) in 1 ml PBS. Cells were incubated sequentially in protein solutions, primary antibody solution (Invitrogen MA1-91878 - monoclonal mouse α- FLAG FG4R), and secondary antibody (R&D systems NL008 - donkey α-mouse IgG conjugated to NorthernLights637) prior to imaging/cytometry. All incubations were 45 min. at 4°C. Samples were processed within 24 h of labeling.

### Microscopy

Protein labeled cells were mounted in 50% SlowFade Gold and visualized with a Zeiss LSM 880 with Airyscan super resolution. Protein binding frequency was quantified manually. For binding intensity, the mean and maximum fluorescent intensity of each cell was quantified in FIJI (53). Episymbiont-host binding assays and twitching motility assays were visualized via Zeiss Axio Widefield microscope. Host binding quantified manually. Cross sectional area quantified in FIJI.

### Flow cytometry

Protein labeled cells were analyzed with an Invitrogen Attune Nxt using the Rl1 channel to detect NorthernLights637 in biological triplicate. All samples were quantified using identical gain settings. On average, ≈395,000 particles were counted per treatment; however, some pilin treated samples had decreased particle counts from potential aggregation (Supplemental table 2). Flow data was analyzed in FlowJo11. Gates were set so that each strain’s unlabeled treatment had approximately 99% fluorescence negative cells.

### Glycan microarray analysis

Purified PilX1 and PilX2 were submitted to Glybrary (Protein IDs 170 and 162). Each was diluted to 5 µg/ml and 50 µg/ml in TSM Binding Buffer (TSMBB; 20 mM tris-HCl, 150 mM NaCl, 2 mM CaCl_2_, 2 mM MgCl_2_, 0.05% Tween 20, 1% BSA, pH 7.4). Samples were applied to glycan microarray chips printed with 313 purified glycans in sextuplicate. Protein labeled chips were rinsed, incubated with mouse α-FLAG antibody conjugated to biotin (Invitrogen MA1-91878-BTIN), rinsed, incubated with Alexa Fluor-488-Streptavidin, rinsed, and imaged via ProScanArray Scanner (Perkin Elmer). Data in Supplemental table 3.

### *N. lyticus* genetics

*N. lyticus* cells were transformed as previously described (5, 6). T4P-1 defective strain generated via antisense knockdown of T4P-1’s major pilin (PilA1; TM7x_00450). Homologous recombination with 300 bp homology arms was used to insert the antisense cassette into the genome at Neutral Site 1 (TM7x NS1::Antisense-*pilA1*). Cassette uses the native *pilA1* promoter to express reverse complement transcripts and the *rpsJ* promoter to express resistance gene HphI. Transformants were selected via serial passaging with 300 μg/ml hygromycin B, then validated via whole genome sequencing (Plasmidsaurus). Primers #22-31 were utilized to generate transformation constructs, primers 32-37 were utilized for knockdown validation (normalized to *gyrA* expression). Primers described in Supplemental table 5.

### Dot blots

Nitrocellulose membranes (0.45 μm) were spotted with *B. subtilis* peptidoglycan (0.1 mg Sigma-Aldrich 69554-10MG-F), whole *S. odontolytica* cell walls (0.1 mg), polysaccharide-depleted cell wall fractions (2 μl at 0.1%), and CWPS fractions (2 ul at 0.1%) in triplicate. PBS was used as vehicle control. Membranes were sequentially blocked in PBST+5% BSA (0.05% tween 20 in PBS), labeled with 50 μg of protein (PilA2, PilX1, or PilX2), labeled with primary antibody (mouse α-FLAG), labeled with secondary antibody (donkey α-mouse NorthernLights637), and imaged on a BioRad ChemiDoc MP imaging system. All incubations were rotated for one h at 4°C. Fluorescent intensity was quantified in FIJI.

### Cell wall and CWPS isolation

Bacterial cell wall purification was based on a peptidoglycan isolation protocol (54) then modified to exclude glycan removal. 500-1500 mL cultures were grown overnight, except for the tunicamycin treatment, which was supplemented with 1 μg/ml tunicamycin and incubated for 48 h. Cells were added into boiling PBS with 8% SDS (w/v) for 30 minutes. Liberated sacculi were treated with DNase (10 μg/mL), RNase (50 μg/mL), and Pronase protease (200 μg/mL). All enzymatic digestions performed at 37°C for 90 minutes. Sacculi were boiled for 30 min. in 1% SDS, rinsed twice in water, rinsed in acetone, then dehydrated. To isolate CWPS, sacculi were suspended in 10% trichloroacetic acid (TCA) at 4°C overnight, centrifuged to collect the supernatant, resuspended in 5% TCA at 97°C for 30 min., and centrifuged to collect the second supernatant (55). Supernatant fractions were combined and dialyzed (3 kD molecular cutoff) prior to desiccation.

### Twitching motility assay

*N. lyticus* motility was measured as previously described (6). Briefly, co-cultures containing *N. lyticus* NS1::HphI, *N. lyticus* NS1::Antisense*pilA1*-HphI, and *N. lyticus* PilA2::HphI were grown to late log phase, transferred to ≈350 μm-thick 0.8% agar pads (w/v), and imaged every 3 seconds for 3 minutes via phase contrast. Heat killed *N. lyticus* NS1::HphI (15 minutes at 97°C) were used as a negative control. Motion paths were traced and quantified in FIJI (53). Each treatment quantified paths for 36 cells, collected from across three distinct replicates/videos. To assess motility as a function of initial host-distance, 63 motion paths were traced per genotype, looking exclusively at cells within 10 μm host-distance.

### *S. odontolytica* binding assay

Prior to re-association experiments, all *N. lyticus* strains had to be isolated as previously described (11). After isolation cells were normalized to an OD_600_ of 0.1. All genotypes had three or more biological replicates. Isolates were re-introduced to *S. odontolytica* hosts at MOI 0.1.

Co-cultures were incubated 24 h, then the percentage of host cells bound by episymbionts was determined via phase contrast microscopy. For every replicate, between 218 and 622 cells (389 average). For exogenous sugar experiments, custom batches of BHI were prepared with sucrose, fucose, mannose, glucose, or rhamnose at 0.2 M, lower concentrations were achieved by diluting with BHI. qPCR primers 32-33 were used to assess quantify episymbionts (Supplemental table 5). For quercetin (100 μM) and tunicamycin (1.2 μM) treatments, 1000x stocks were prepared fresh in DMSO before use. Nanosynbacteria competition experiments were performed in biological triplicate. For tripartite competition, *N. lyticus*, BB004, and TM7-008 were introduced into *S. odontolytica* cultures with individual MOIs of 0.1, followed by outgrowth and passaging (with or without 50 mM L-rhamnose) every 24 h for 12 days.

### *S. odontolytica* genetics

*S. odontolytica* XH001 was made electrocompetent and transformed as previously described (32, 56, 57). Briefly, to generate gene knockout constructs 1,000 bp homology arms flanking *marR* (APY09_RS07555), the FucoseLocus region (APY09_RS05345 - APY09_RS05355), *rmlC-D* (APY09_RS00100), *rgpB* (APY09_RS05560), *rgpF* (APY09_RS05510), *tagO* (APY09_RS06605), a putative galactosyltransferase (APY09_RS05455), a putative N-acetyl glucosamine transferase/*gacI* (APY09_RS00090), and a convergently encoded neutral site termed NSXH (between APY09_RS05760 and APY09_RS05765) were amplified and assembled into allelic exchange constructs using either the Kanamycin resistance cassette from pJRD215 (58), the chloramphenicol resistance cassette from pCWU6 (59), or the Erythromycin resistance cassette from pVA2198 (60). Primers 38-103 were used to construct the above deletion constructs and primers 104-121 were used for validation (Supplemental table 5). Electroporation was performed with 1000-2000 ng of template at 25 mF, 2.5 kV, and 400 ohms. Selection utilized 400 μg/mL kanamycin, 10 μg/mL thiamphenicol, or 300 μg/mL erythromycin as appropriate.

### Modified anthrone assay

Our modified anthrone assay was based on previous analyses of Rha-CWPS in *Streptococcus mutans* (33). Briefly, a 0.2% (w/v) solution of anthrone in concentrated sulfuric acid was prepared. 1 mL of anthrone solution was added to 200 μL of monosaccharides (0.1% and 0.01% w/v) or bacterial samples (2 mg and 0.2 mg mass) suspended in water. Samples were heated to 97°C for 10 min. prior to spectroscopy. Optical densities were measured at 580 nm (representing all carbohydrates present) and 700 nm (representing only non-6DH saccharides). 6DH:non-6DH ratio was calculated as (OD_580_-OD_700_)/OD_700_.

### HPLC analysis of polysaccharides

Fluorescent labelling of CWPS fractions was conducted as described previously (61). Briefly, 50 μL of 1 mg/mL CWPS solution was mixed with 25 μL of freshly prepared label (48 mg/mL 2-aminobenzamide (Ludger) in DMSO/acetic acid (85:15 v/v)). Then, 25 μL of 1 M 1-picolane-borane in DMSO was added to achieve a volume of 100 μL. The reaction was incubated at 65 °C for 2 h then centrifuged to remove precipitates. Labelled samples were separated by size-exclusion chromatography on a Phenomenex BioZen 1.8 µM SEC-2 column (300 × 4.6 mm, 00H-4769-E0) at room temperature, using a Dionex Ultimate 3000 uHPLC system operated by Chromeleon Chromatography Data System (v7.2 SR4) (62). The mobile phase consisted of 0.1 M phosphate buffer (pH 6.8) with 0.025% sodium azide, run isocratically at a flow rate of 0.400 mL/min for 30 min. Labelled CWPS were detected via fluorescence (excitation 320 nm, emission 420 nm).

### Polysaccharide acid hydrolysis and HPAED-PAD analysis

1 mg of dried CWPS was resuspended in 1 ml of 300 mM HCl and incubated 1 h at 100 °C. The sample was neutralized with 1 M NaOH. Hydrolysate samples were dried under a nitrogen stream. Samples were analyzed using Ion Chromatography with Pulsed Amperometric Detection (IC-PAD) using an ICS-6000 system equipped with a CARBOPAC PA1 anion exchange column (ThermoFisher). Detection was facilitated by a gold working electrode and a PdH reference electrode, using a Carbo Quad waveform. Buffer A consisted of 100 mM NaOH and Buffer B was 100 mM NaOH and 0.5 M sodium acetate. Each sample was run at a constant flow rate of 0.25 mL/min for 40 min using the following method: 0–20 min, isocratic elution with 100% Buffer A; 20–25 min, isocratic elution with 100% Buffer B. Data were processed using Chromeleon™ Chromatography Management System V.6.8.

### Quantification and Statistical analysis

All statistical analyses performed in Prism 10.3.0 or 10.6.1 (GraphPad Software, San Diego, CA). Most statistical comparisons were analyzed via one-way ANOVA with Dunnett’s multiple comparison test. For proximity induced motility assays, logistical regressions, or stepwise logistical regressions, were used to determine likelihood of motility (defined as >5 μm/min). qPCR quantification equations derived via semi-log regression of ct values for serial dilution series with known concentrations.

## Materials availability

All unique strains/reagents generated in this study are available upon request but may require completion of a materials transfer agreement. Requests for further information and resources should be directed to and will be fulfilled by the lead contact, Dr. Batbileg Bor

## Data and code availability

Genomic data for organisms utilized within this study are present within NCBI or GTDB. Protein samples applied to glycan microarrays are described in the Glybrary database (Protein IDs 162 and 170). This study contains discussion of published transcriptomic data (7) available at GEO GSE196744. This paper does not report original code.

## Supporting information

Supplemental table 1: DALI structural similarity analysis results

Supplemental table 2: Summary flow cytometry statistics

Supplemental table 3: Glycan microarray results

Supplemental table 4: Compiled list of all strains and their genotypes

Supplemental table 5: Compiled list of all primers used in this study

Supplemental video 1: Examples of N. lyticus twitching motility with and without T4P function

## Acknowledgements

We thank Dr. Larry Gallagher for experiments and discussions that informed the selection of target genes with the host. This research was partially supported by grants from the National Institute of Dental and Craniofacial Research under Awards 1R01DE031274 (BB), 1R01DE023810 (XH, BB), and T90 DE026110-07 (ASG). This work utilized a Zeiss LSM880 and an AXIO widefield microscope housed in the ADA Forsyth Institute Advanced Microscopy Core Facility (RRID:SCR_021121) supported by NIH grant 1S10OD034405-01. KA was funded by a studentship from the Darwin Trust of Edinburgh. JLR was supported by the Wellcome Trust through Investigator Award 226644/Z/22/Z.

## Authors’ contributions

Conceptualization by ASG, PJM, JDM, and BB. Methodology by ASG, JGSS, LL, KA, MX, and BB. Software by ASG and YW. Formal analysis by ASG, KA, YW, JLR, and PJM. Investigation by ASG, JGSS, LL, KA, MX, YW, and JLR. Resources provided by ASG, ECL, XH, PJM, JDM, and BB. Data Curation by ASG and BB. Original Draft written by ASG. Review and Editing by all authors. Visualization by ASG. Supervision by PJM, JDM, and BB. Funding acquisition by ECL, XH, PJM, JDM, and BB.

---

**Supplemental figure 1.**
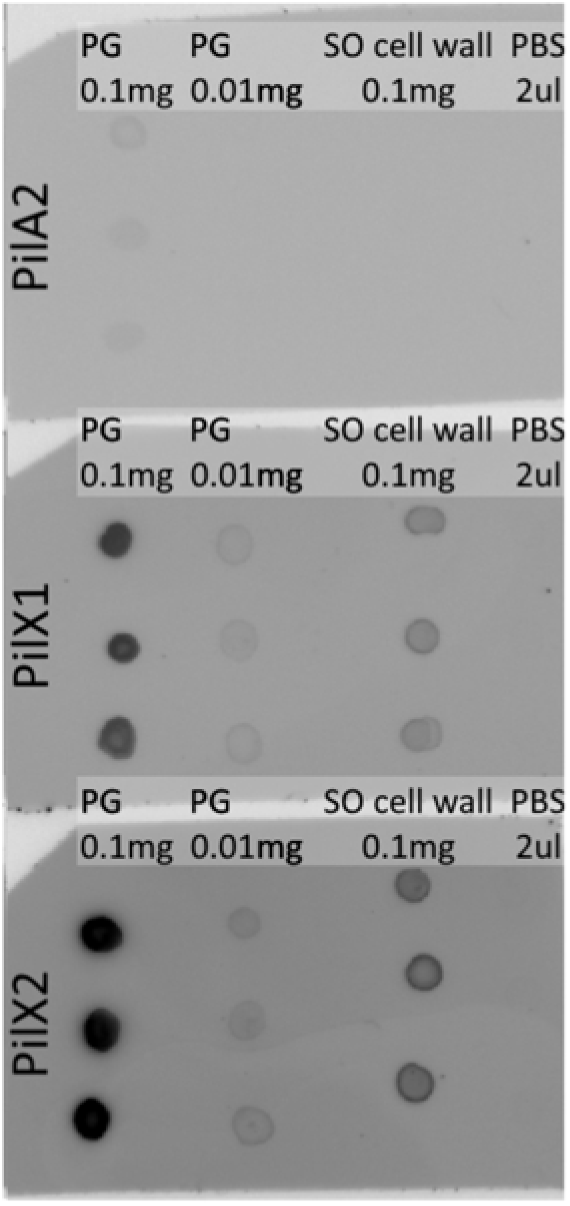
PilX1 and PilX2 bind pure peptidoglycan better than host-derived cell walls. A representative immuno-dot blot replicate showing PilA2, PilX1, and PilX2 binding of pure peptidoglycan (derived from *B. subtilis*) and whole cell wall isolates from *S. odontolytica*. Samples were thoroughly sonicated prior to spotting to ensure small particle sizes.

**Supplemental figure 2.**
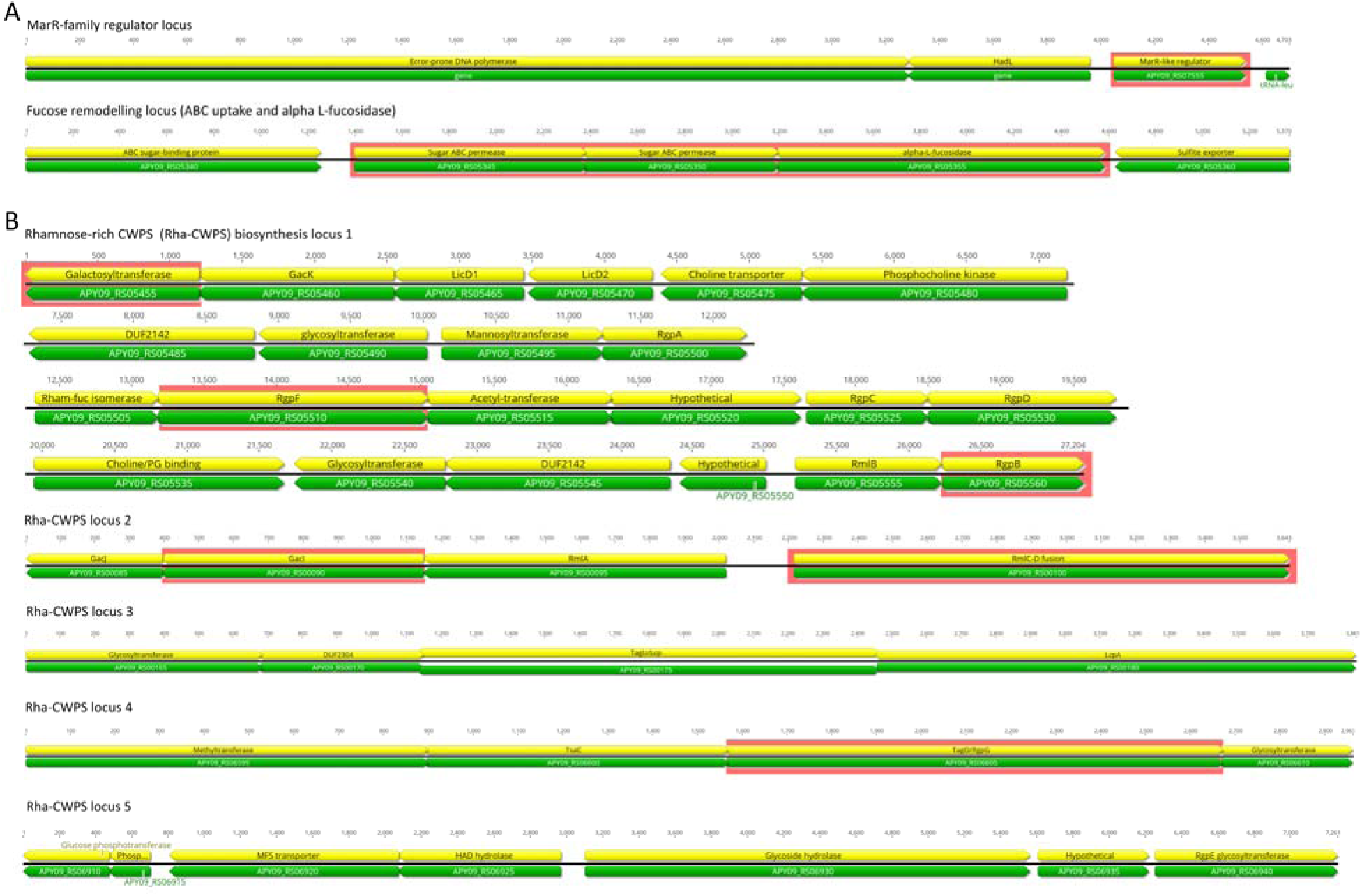
Genomic loci encoding CWPS biosynthesis genes. **(A)** *marR* has no obvious signs of being encoded within an operon but is located near highly conserved genes like DNA polymerase III and a tRNA-leu gene. The fucose remodeling locus contains genes predicted to be involved in processing and uptake of fucose. These include ABC sugar binding proteins, an ABC permease, and an L-fucosidase. We deleted three of these genes, APY_RS05345-APY_RS05355, which overlap and are likely expressed as an operon. **(B)** Genomic investigation revealed the presence of rhamnose-rich CWPS biosynthesis loci homologous to those found in *Streptococcus* species. All genes/operons that were targeted for knockout have red highlighting; however, all Rha-CWPS genes except for *gacI* were essential for survival.

**Supplemental figure 3.**
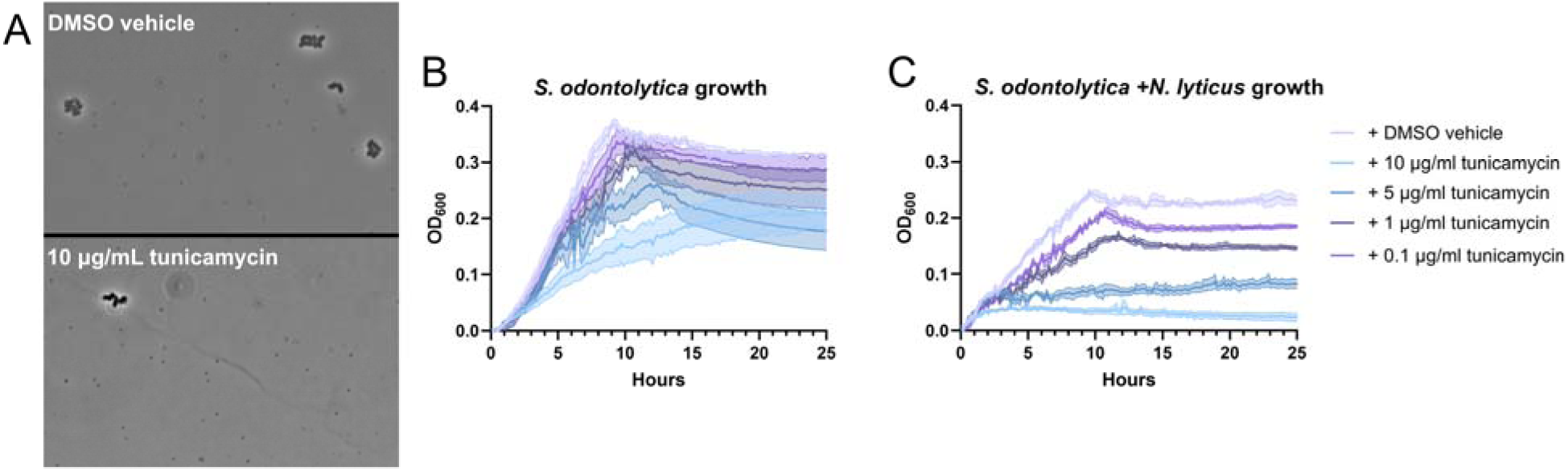
Optimization of tunicamycin treatment. Since most Rha-CWPS genes were determined to be essential for survival, a chemical inhibitor of the CWPS synthesis protein TagO called tunicamycin was used to disrupt synthesis. **(A)** Tunicamycin did not appear to inhibit *N. lyticus* cells, as episymbionts were plentiful at all assayed concentrations. **(B)** However, Tunicamycin did inhibit *S. odontolytica* growth at all assayed concentrations, when monitored via growth curve. **(C)** Tunicamycin inhibited *S. odontolytica* synergistically with *N. lyticus* when grown in co-culture, and at 10 μg/ml these combined stressors completely halted growth. Based on these results, tunicamycin treatments were performed at 0.1 μg/ml to minimize host cell stress.

**Supplemental figure 4.**
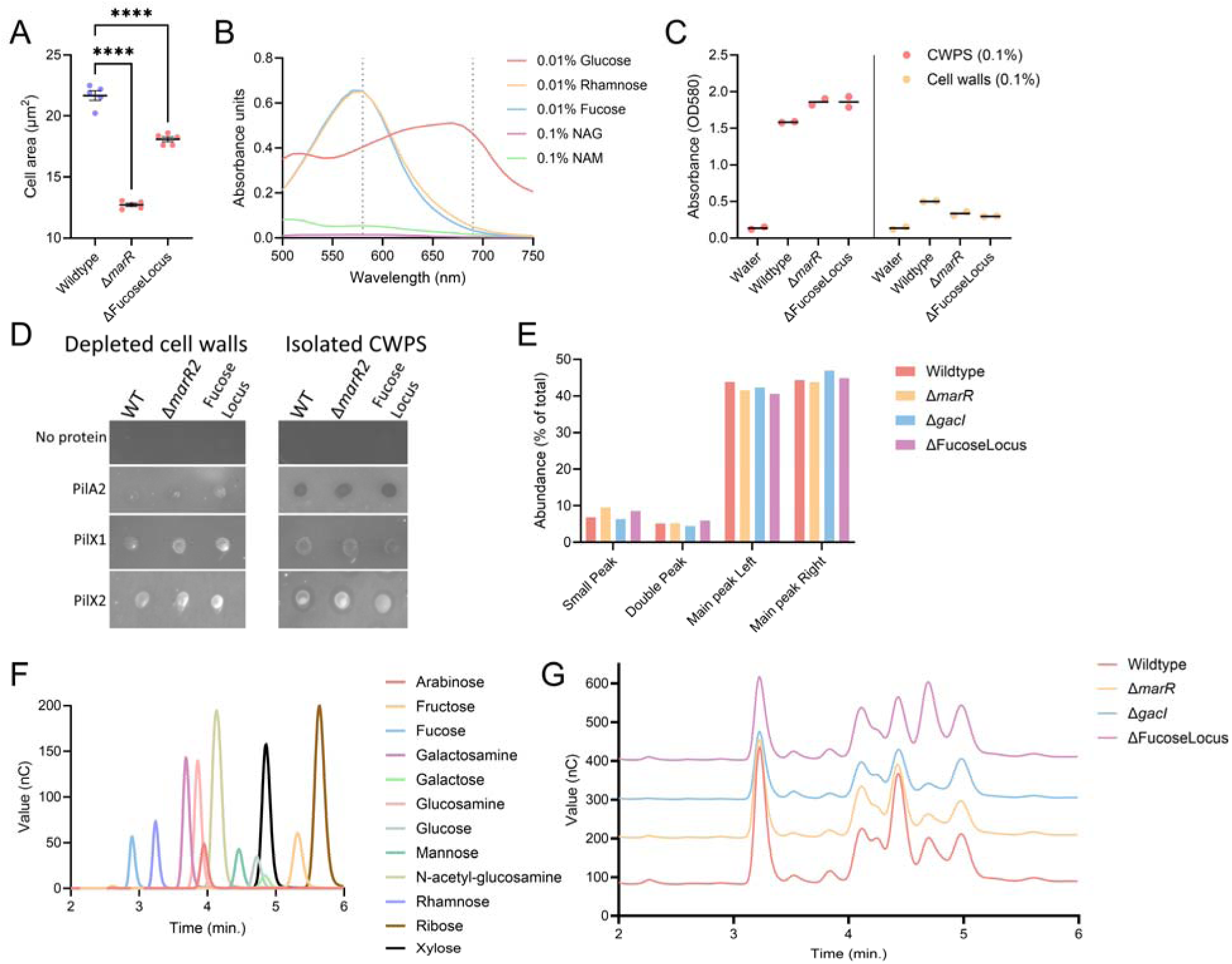
Analysis of CWPS biosynthesis mutants. **(A)** *S. odontolytica* Δ*marR* and *S. odontolytica* ΔFucoseLocus cells were significantly shorter on average than wildtype cells. Since this organism can have pleomorphic shapes, cross-sectional cell area was used instead of cell length. **(B)** The anthrone assay hydrolyzes saccharides into colorimetric products which can be differentiated by absorbance pattern. These monosaccharide standards display how 6-deoxyhexoses generate a product with a relatively narrow absorbance peak around 580 nm, while glucose produces a product with a wider absorbance peak. N-acetylglucosamine (NAG) and N-acetylmuramic acid (NAM) from peptidoglycan contribute little to no product from this reaction. **(C)** The anthrone assay confirms that TCA cleavage liberates the majority of CWPS from cell walls, though some residual saccharides remain attached, particularly in wildtype cells. **(D)** An example replicate of the dot blots prepared with glycan depleted cell walls and the isolated CWPS fractions. **(E)** Using HPLC to separate glycans within collected CWPS fractions, five peaks were detected in all samples. Relative abundances were similar between all treatments. **(F)** When performing HPAEC-PAD, we developed a set of monosaccharide standards common in known Gram-positive CWPS. **(G)** These standards were compared to the peaks present in CWPS hydrolysates to quantify the relative abundance of each component, here shown with representative traces.

**Supplemental figure 5.**
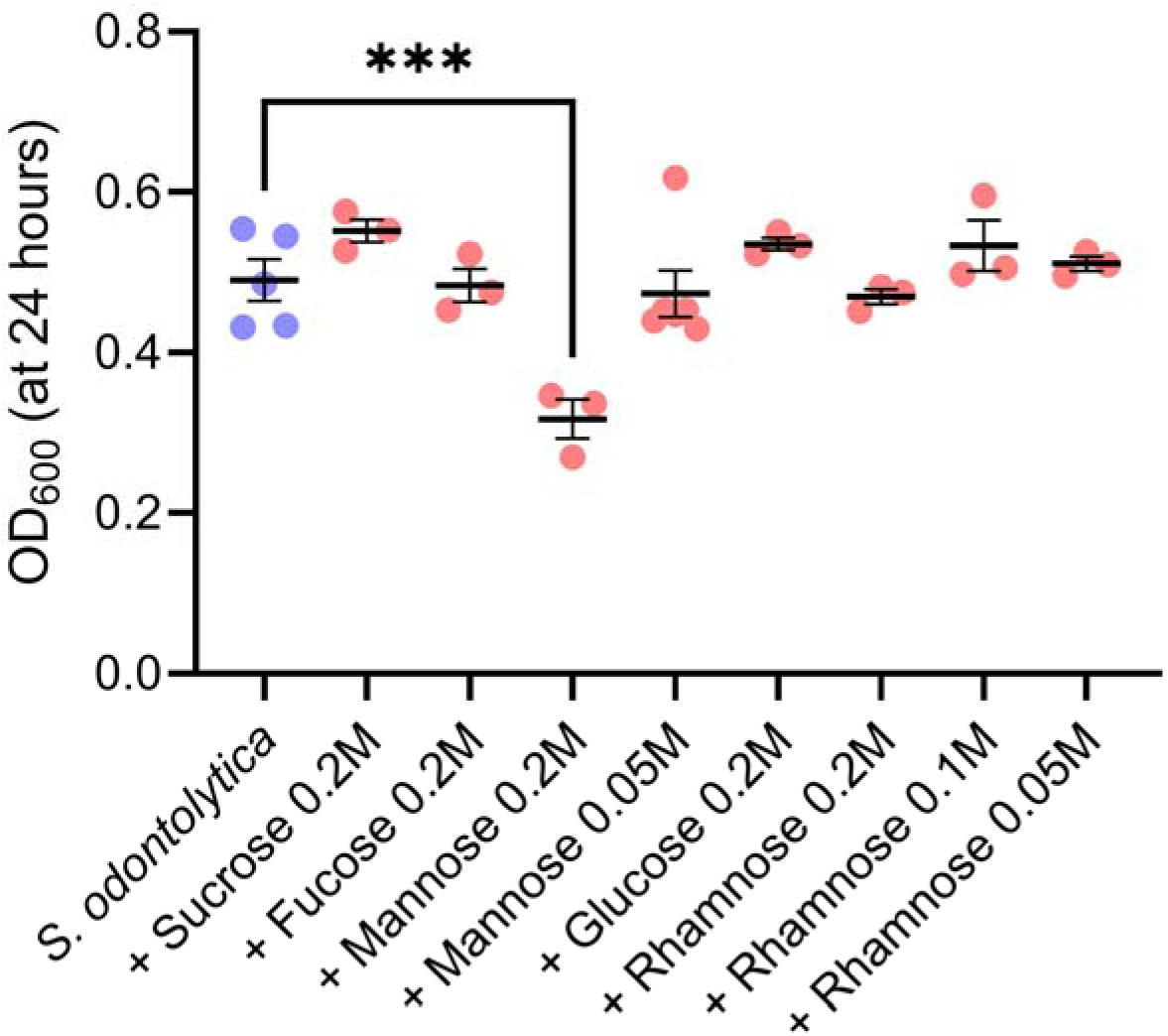
Exogenous mannose induces sugar toxicity in *S. odontolytica*. Cultivation of *S. odontolytica* monocultures within medium supplemented with monosaccharides known to occur in Gram-positive CWPS revealed that there was no solute toxicity at 0.2 M concentrations, however high concentrations of mannose specifically inhibited growth.

## Supplemental Items

Supplemental table 1: DALI structural similarity analysis results

Supplemental table 2: Summary flow cytometry statistics

Supplemental table 3: Glycan microarray results

Supplemental table 4: Compiled list of all strains and their genotypes

Supplemental table 5: Compiled list of all primers used in this study

Supplemental video 1: Examples of *N. lyticus* twitching motility with and without T4P function

## References

1. Méheust R, Burstein D, Castelle CJ, Banfield JF. 2019. The distinction of CPR bacteria from other bacteria based on protein family content. Nat Commun 10:4173.

2. Castelle CJ, Banfield JF. 2018. Major new microbial groups expand diversity and alter our understanding of the tree of life. Cell 172:1181–1197.

3. Brown CT, Hug LA, Thomas BC, Sharon I, Castelle CJ, Singh A, Wilkins MJ, Wrighton KC, Williams KH, Banfield JF. 2015. Unusual biology across a group comprising more than 15% of domain Bacteria. Nature 523:208–211.

4. Hug LA, Baker BJ, Anantharaman K, Brown CT, Probst AJ, Castelle CJ, Butterfield CN, Hernsdorf AW, Amano Y, Ise K, Suzuki Y, Dudek N, Relman DA, Finstad KM, Amundson R, Thomas BC, Banfield JF. 2016. A new view of the tree of life. Nat Microbiol 1:16048.

5. Wang Y, Gallagher LA, Andrade PA, Liu A, Humphreys IR, Turkarslan S, Cutler KJ, Arrieta-Ortiz ML, Li Y, Radey MC, McLean JS, Cong Q, Baker D, Baliga NS, Peterson SB, Mougous JD. 2023. Genetic manipulation of Patescibacteria provides mechanistic insights into microbial dark matter and the epibiotic lifestyle. Cell 186:4803–4817.e13.

6. Grossman AS, Lei L, Botting JM, Liu J, Nahar N, Liu J, McLean JS, He X, Bor B. 2025. Saccharibacteria deploy two distinct type IV pili, driving episymbiosis, host competition, and twitching motility. ISME J wraf119.

7. Hendrickson EL, Bor B, Kerns KA, Lamont EI, Chang Y, Liu J, Cen L, Schulte F, Hardt M, Shi W, He X, McLean JS. 2022. Transcriptome of epibiont Saccharibacteria *Nanosynbacter lyticus* strain TM7x during the establishment of symbiosis. J Bacteriol 204:e00112–22.

8. Xie B, Wang J, Nie Y, Tian J, Wang Z, Chen D, Hu B, Wu X-L, Du W. 2022. Type IV pili trigger episymbiotic association of Saccharibacteria with its bacterial host. Proceedings of the National Academy of Sciences 119:e2215990119.

9. McLean JS, Bor B, Kerns KA, Liu Q, To TT, Solden L, Hendrickson EL, Wrighton K, Shi W, He X. 2020. Acquisition and adaptation of ultra-small parasitic reduced genome bacteria to mammalian hosts. Cell Rep 32:107939.

10. He X, McLean JS, Edlund A, Yooseph S, Hall AP, Liu S-Y, Dorrestein PC, Esquenazi E, Hunter RC, Cheng G, Nelson KE, Lux R, Shi W. 2015. Cultivation of a human-associated TM7 phylotype reveals a reduced genome and epibiotic parasitic lifestyle. Proceedings of the National Academy of Sciences 112:244–249.

11. Bor B, McLean JS, Foster KR, Cen L, To TT, Serrato-Guillen A, Dewhirst FE, Shi W, He X. 2018. Rapid evolution of decreased host susceptibility drives a stable relationship between ultrasmall parasite TM7x and its bacterial host. Proceedings of the National Academy of Sciences 115:12277–12282.

12. Wang Y, Colautti J, Gallagher LA, Lee J, Tan Y, Farrell S, Farrell C, Bor B, Whitney JC, Zhang D, Peterson SB, Mougous JD. 2026. Oral Actinobacteria sense and defend against parasitic epibionts. bioRxiv 2026.09.08.750141.

13. Rosan B, Hammond BF. 1974. Extracellular Polysaccharides of *Actinomyces viscosus*. Infect Immun 10:304–308.

14. Mistou M-Y, Sutcliffe IC, van Sorge NM. 2016. Bacterial glycobiology: rhamnose-containing cell wall polysaccharides in Gram-positive bacteria. FEMS Microbiol Rev 40:464–479.

15. Zaychikov VA, Potekhina N V, Dmitrenok AS, Fan D, Tul’skaya EM, Dorofeeva L V, Evtushenko LI. 2021. Cell wall rhamnan in Actinobacteria of the genus *Curtobacterium*. Microbiology (N Y) 90:343–348.

16. Shashkov AS, Tul’skaya EM, Streshinskaya GM, Dmitrenok AS, Potekhina N V, Senchenkova SN, Piskunkova NF, Dorofeeva L V, Evtushenko LI. 2020. Rhamnomannans and teichuronic acid from the cell wall of *Rathayibacter tritici* VKM Ac-1603T. Biochemistry (Moscow) 85:369–377.

17. Shashkov AS, Potekhina N V, Tul’skaya EM, Dmitrenok AS, Senchenkova SN, Dorofeeva L V, Evtushenko LI. 2024. Novel rhamnose-containing glycopolymers from the cell wall of *Rathayibacter festucae* VKM Ac-1390T. Biochemistry (Moscow) 89:2204–2213.

18. Kovacs CJ, Faustoferri RC, Bischer AP, Quivey Jr. RG. 2019. *Streptococcus mutans* requires mature rhamnose-glucose polysaccharides for proper pathophysiology, morphogenesis, and cellular division. Mol Microbiol 112:944–959.

19. Guérin H, Courtin P, Guillot A, Péchoux C, Mahony J, van Sinderen D, Kulakauskas S, Cambillau C, Touzé T, Chapot-Chartier M-P. 2024. Molecular mechanisms underlying the structural diversity of rhamnose-rich cell wall polysaccharides in lactococci. Journal of Biological Chemistry 300.

20. Guérin H, Kulakauskas S, Chapot-Chartier M-P. 2022. Structural variations and roles of rhamnose-rich cell wall polysaccharides in Gram-positive bacteria. Journal of Biological Chemistry 298.

21. Varadi M, Bertoni D, Magana P, Paramval U, Pidruchna I, Radhakrishnan M, Tsenkov M, Nair S, Mirdita M, Yeo J, Kovalevskiy O, Tunyasuvunakool K, Laydon A, Žídek A, Tomlinson H, Hariharan D, Abrahamson J, Green T, Jumper J, Birney E, Steinegger M, Hassabis D, Velankar S. 2024. AlphaFold Protein Structure Database in 2024: Providing structure coverage for over 214 million protein sequences. Nucleic Acids Res 52:D368– D375.

22. Holm L, Laiho A, Törönen P, Salgado M. 2023. DALI shines a light on remote homologs: One hundred discoveries. Protein Science 32:e4519.

23. Holm L. 2022. Dali server: structural unification of protein families. Nucleic Acids Res 50:W210–W215.

24. Johnson MDL, Garrett CK, Bond JE, Coggan KA, Wolfgang MC, Redinbo MR. 2011. *Pseudomonas aeruginosa* PilY1 binds Integrin in an RGD- and calcium-dependent manner. PLoS One 6:e29629-.

25. Vinogradov E V, Krajewka-Pietrasik D, Kaca W, Shashkov AS, Knirel YA, Kochetkov NK. 1989. Structure of *Proteus mirabilis* 027 O-specific polysaccharide containing amino acids and phosphoethanolamine. Eur J Biochem 185:645–650.

26. Senchenkova SN, Perepelov A V, Cedzynski M, Swierzko AS, Ziolkowski A, Shashkov AS, Kaca W, Knirel YA, Jansson P-E. 2004. Structure of a highly phosphorylated O-polysaccharide of *Proteus mirabilis* O41. Carbohydr Res 339:1347–1352.

27. Pradip A, Gwen A, Belinda W, K VN. 1999. Serotype 1a O-Antigen Modification: Molecular Characterization of the Genes Involved and Their Novel Organization in the *Shigella flexneri* Chromosome. J Bacteriol 181:4711–4718.

28. Lu F, N SS, Jinghua Y, S SA, Jiang T, Hongjie G, Guang Z, A KY, Peter R, Lei W. 2004. Structural and Genetic Characterization of the *Shigella boydii* Type 13 O Antigen. J Bacteriol 186:383–392.

29. Irina S, Evgeny V, Pascal C, Julija A, Mickael M, Efstathios G, Simon P, Sylviane F, Christine P, Stuart A, Jennifer M, Douwe van S, Saulius K, Yann G, Marie-Pierre C-C. 2017. Another brick in the wall: A rhamnan polysaccharide trapped inside peptidoglycan of *Lactococcus lactis*. mBio 8:10.1128/mbio.01303-17.

30. Gisch N, Kohler T, Ulmer AJ, Müthing J, Pribyl T, Fischer K, Lindner B, Hammerschmidt S, Zähringer U. 2013. Structural Reevaluation of *Streptococcus pneumoniae* Lipoteichoic Acid and New Insights into Its Immunostimulatory Potency. Journal of Biological Chemistry 288:15654–15667.

31. Mücke P-A, Ostrzinski A, Hammerschmidt S, Maaß S, Becher D. 2020. Proteomic Adaptation of *Streptococcus pneumoniae* to the Antimicrobial Peptide Human Beta Defensin 3 (hBD3) in Comparison to Other Cell Surface Stresses. Microorganisms 8.

32. Bedree JK, Bourgeois J, Balani P, Cen L, Hendrickson EL, Kerns KA, Camilli A, McLean JS, Shi W, He X. 2024. Identifying essential genes in *Schaalia odontolytica* using a highly-saturated transposon library. bioRxiv 2024.07.17.604004.

33. Zamakhaeva S, Chaton CT, Rush JS, Ajay Castro S, Kenner CW, Yarawsky AE, Herr AB, van Sorge NM, Dorfmueller HC, Frolenkov GI, Korotkov K V, Korotkova N. 2021. Modification of cell wall polysaccharide guides cell division in *Streptococcus mutans*. Nat Chem Biol 17:878–887.

34. Potekhina NV, Shashkov AS, Evtushenko LI, Gavrish EY, Senchenkova SN, Stomakhin AA, Usov AI, Naumova IB, Stackebrandt E. 2003. A novel mannitol teichoic acid with side phosphate groups of Brevibacterium sp. VKM Ac-2118. Eur J Biochem 270:4420–4425.

35. Boulanger EF, Sabag-Daigle A, Thirugnanasambantham P, Gopalan V, Ahmer BM. 2021. Sugar-phosphate toxicities. Microbiology and Molecular Biology Reviews 85:e00123–21.

36. Jie N, Utter DR, Kerns KA, Lamont IE, Hendrickson EL, Liu J, Wu T, He X, McLean J, Bor B. 2022. Strain-level variation and diverse host bacterial responses in episymbiotic Saccharibacteria. mSystems 7:e01488–21.

37. Chipashvili O, Utter DR, Bedree JK, Ma Y, Schulte F, Mascarin G, Alayyoubi Y, Chouhan D, Hardt M, Bidlack F, Hasturk H, He X, McLean JS, Bor B. 2021. Episymbiotic Saccharibacteria suppresses gingival inflammation and bone loss in mice through host bacterial modulation. Cell Host Microbe 29:1649–1662.e7.

38. Grossman AS, Weng J, Silverman AD, Bor B. 2026. Resolving host-episymbiont interaction dynamics through continuous cultivation. mSystems 0:e00592–26.

39. Lechevalier MP, Lechevalier H. 1970. Chemical composition as a criterion in the classification of aerobic actinomycetes. International Journal of Systemic Bacteriology 20:435–443.

40. Cummins CS, Harris H. 1958. Studies on the Cell-Wall Composition and Taxonomy of *Actinomycetales* and Related Groups. Microbiology (N Y) 18:173–189.

41. van den Berg B, Silale A, Baslé A, Brandner AF, Mader SL, Khalid S. 2022. Structural basis for host recognition and superinfection exclusion by bacteriophage T5. Proceedings of the National Academy of Sciences 119:e2211672119.

42. Bondy-Denomy J, Qian J, Westra ER, Buckling A, Guttman DS, Davidson AR, Maxwell KL. 2016. Prophages mediate defense against phage infection through diverse mechanisms. ISME J 10:2854–2866.

43. Kumar A, Sýkorová P, Demo G, Dobeš P, Hyršl P, Wimmerová M. 2016. A novel fucose-binding lectin from *Photorhabdus luminescens* (PLL) with an unusual heptabladed beta-propeller tetrameric structure. Journal of Biological Chemistry 291:25032–25049.

44. Teufel F, Almagro Armenteros JJ, Johansen AR, Gíslason MH, Pihl SI, Tsirigos KD, Winther O, Brunak S, von Heijne G, Nielsen H. 2022. SignalP 6.0 predicts all five types of signal peptides using protein language models. Nat Biotechnol 40:1023–1025.

45. Abramson J, Adler J, Dunger J, Evans R, Green T, Pritzel A, Ronneberger O, Willmore L, Ballard AJ, Bambrick J, Bodenstein SW, Evans DA, Hung C-C, O’Neill M, Reiman D, Tunyasuvunakool K, Wu Z, Žemgulytė A, Arvaniti E, Beattie C, Bertolli O, Bridgland A, Cherepanov A, Congreve M, Cowen-Rivers AI, Cowie A, Figurnov M, Fuchs FB, Gladman H, Jain R, Khan YA, Low CMR, Perlin K, Potapenko A, Savy P, Singh S, Stecula A, Thillaisundaram A, Tong C, Yakneen S, Zhong ED, Zielinski M, Žídek A, Bapst V, Kohli P, Jaderberg M, Hassabis D, Jumper JM. 2024. Accurate structure prediction of biomolecular interactions with AlphaFold 3. Nature 630:493–500.

46. Schrodinger LLC. 2015. The PyMOL Molecular Graphics System, Version 1.8.

47. Ye Y, Godzik A. 2003. Flexible structure alignment by chaining aligned fragment pairs allowing twists. Bioinformatics 19:ii246–ii255.

48. Campos BM, Liberato MV, Alvarez TM, Zanphorlin LM, Ematsu GC, Barud H, Polikarpov I, Ruller R, Gilbert HJ, Zeri AC de M, Squina FM. 2016. A novel carbohydrate-binding module from sugar cane soil metagenome featuring unique structural and carbohydrate affinity properties. Journal of Biological Chemistry 291:23734–23743.

49. Sýkorová P, Novotná J, Demo G, Pompidor G, Dubská E, Komárek J, Fujdiarová E, Houser J, Hároníková L, Varrot A, Shilova N, Imberty A, Bovin N, Pokorná M, Wimmerová M. 2020. Characterization of novel lectins from *Burkholderia pseudomallei* and *Chromobacterium violaceum* with seven-bladed β-propeller fold. Int J Biol Macromol 152:1113–1124.

50. Bittrich S, Segura J, Duarte JM, Burley SK, Rose Y. 2024. RCSB protein data bank: Exploring protein 3D similarities via comprehensive structural alignments. Bioinformatics 40:btae370.

51. Wang J, Chitsaz F, Derbyshire MK, Gonzales NR, Gwadz M, Lu S, Marchler GH, Song JS, Thanki N, Yamashita RA, Yang M, Zhang D, Zheng C, Lanczycki CJ, Marchler-Bauer A. 2023. The conserved domain database in 2023. Nucleic Acids Res 51:D384–D388.

52. Singh SM, Panda AK. 2005. Solubilization and refolding of bacterial inclusion body proteins. J Biosci Bioeng 99:303–310.

53. Schindelin J, Arganda-Carreras I, Frise E, Kaynig V, Longair M, Pietzsch T, Preibisch S, Rueden C, Saalfeld S, Schmid B, Tinevez J-Y, White DJ, Hartenstein V, Eliceiri K, Tomancak P, Cardona A. 2012. Fiji: An open-source platform for biological-image analysis. Nat Methods 9:676–682.

54. Priyadarshini R, Popham DL, Young KD. 2006. Daughter cell separation by penicillin-binding proteins and peptidoglycan amidases in *Escherichia coli*. J Bacteriol 188:5345– 5355.

55. Potekhina NV, Shashkov AS, Evtushenko LI, Gavrish EY, Senchenkova SN, Stomakhin AA, Usov AI, Naumova IB, Stackebrandt E. 2003. A novel mannitol teichoic acid with side phosphate groups of Brevibacterium sp. VKM Ac-2118. Eur J Biochem 270:4420–4425.

56. Bedree JK, Bor B, Cen L, Edlund A, Lux R, McLean JS, Shi W, He X. 2018. Quorum sensing modulates the epibiotic-parasitic relationship between Actinomyces odontolyticus and its Saccharibacteria epibiont, a Nanosynbacter lyticus Strain, TM7x. Front Microbiol 9.

57. Wu C, Huang I-H, Chang C, Reardon-Robinson ME, Das A, Ton-That H. 2014. Lethality of sortase depletion in *Actinomyces oris* caused by excessive membrane accumulation of a surface glycoprotein. Mol Microbiol 94:1227–1241.

58. Morona R, Mavris M, Fallarino A, Manning PA. 1994. Characterization of the rfc region of *Shigella flexneri*. J Bacteriol 176:733–747.

59. Bibek GC, Peng Z, Arindam N, Jianhua G, Chenggang W. 2023. Development of a xylose-inducible promoter and riboswitch combination system for manipulating gene expression in *Fusobacterium nucleatum*. Appl Environ Microbiol 89:e00667–23.

60. Masayuki T, Thonthi K, Margaret D, Nobushiro H, Howard K. 1998. Role of Arg-gingipain A in virulence of *Porphyromonas gingivalis*. Infect Immun 66:1159–1166.

61. Ruhaak LR, Steenvoorden E, Koeleman CAM, Deelder AM, Wuhrer M. 2010. 2-Picoline-borane: A non-toxic reducing agent for oligosaccharide labeling by reductive amination. Proteomics 10:2330–2336.

62. Franklin A, Salgueiro VC, Layton AJ, Sullivan R, Mize T, Vázquez-Iniesta L, Benedict ST, Gurcha SS, Anso I, Besra GS, Banzhaf M, Lovering AL, Williams SJ, Guerin ME, Scott NE, Prados-Rosales R, Lowe EC, Moynihan PJ. 2024. The mycobacterial glycoside hydrolase LamH enables capsular arabinomannan release and stimulates growth. Nat Commun 15:5740.

