## Supplemental figures 1-5 for "Host Cell Wall Polysaccharides Modulate Nanosynbacteria Recognition and Attachment"

**Supplemental Figures for Host Cell Wall Polysaccharides Modulate Nanosynbacteria Recognition and Attachment**

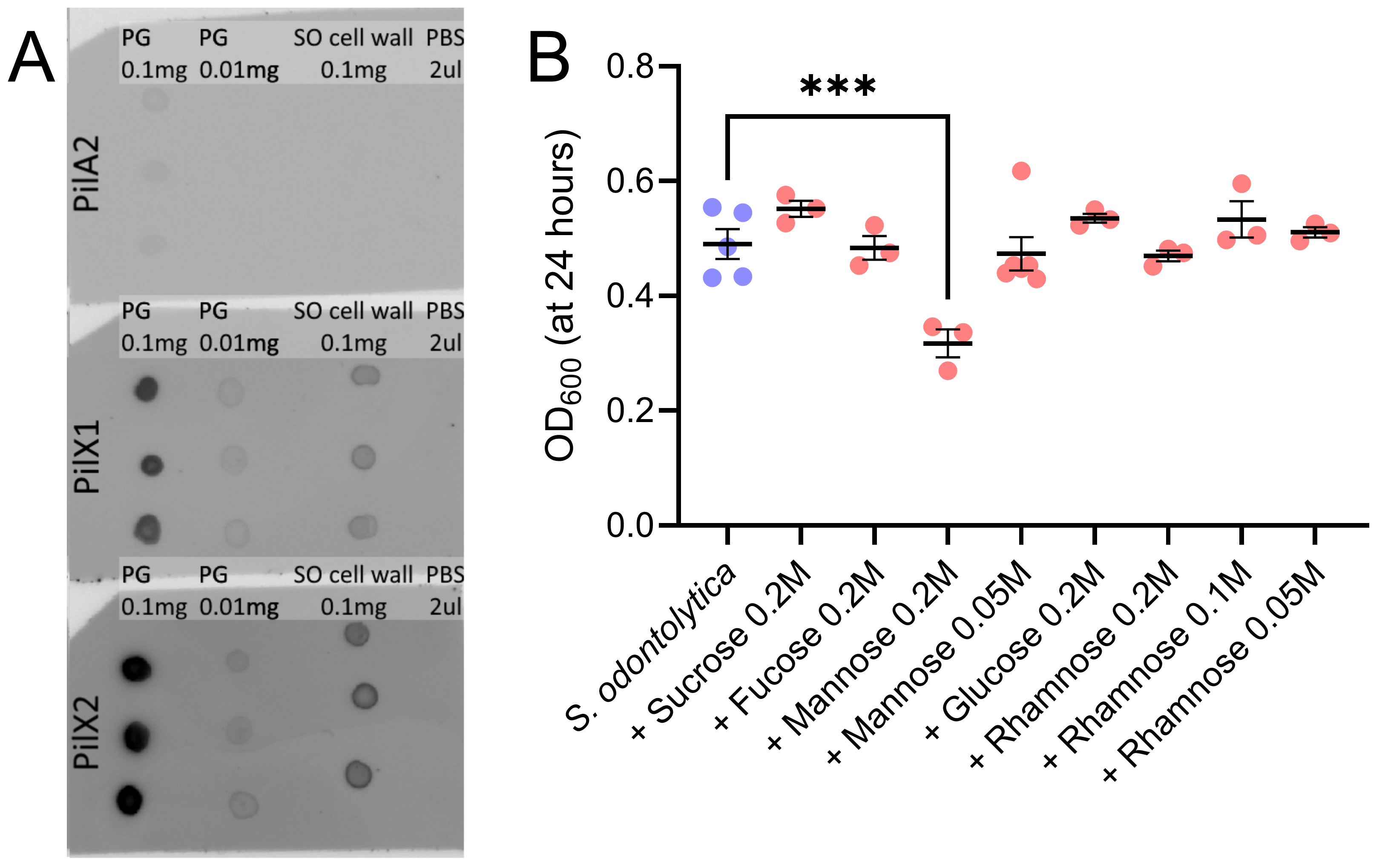

**Supplemental figure 1. PilX1 and PilX2 bind pure peptidoglycan better than host-derived cell walls.** A representative immuno-dot blot replicate showing PilA2, PilX1, and PilX2 binding of pure peptidoglycan (derived from *B. subtilis*) and whole cell wall isolates from *S. odontolytica*. Samples were thoroughly sonicated prior to spotting to ensure small particle sizes.

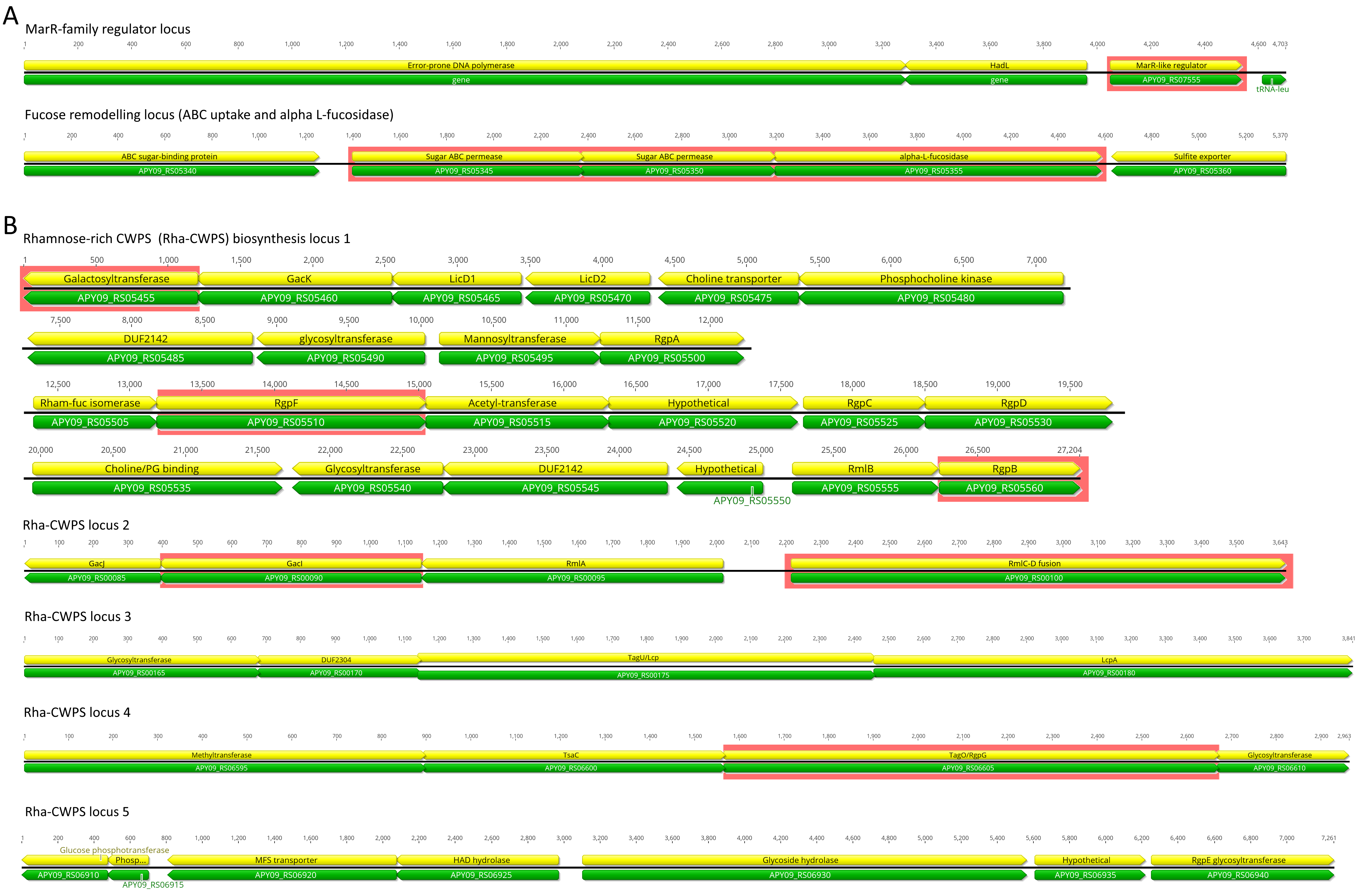

**Supplemental figure 2. Genomic loci encoding CWPS biosynthesis genes. (A)** *marR* has no obvious signs of being encoded within an operon but is located near highly conserved genes like DNA polymerase III and a tRNA-leu gene. The fucose locus contains genes predicted to be involved in processing and uptake of fucose. These include ABC sugar binding proteins, an ABC permease, and an L-fucosidase. We deleted three of these genes, APY_RS05345-APY_RS05355, which overlap and are likely expressed as an operon. **(B)** Genomic investigation revealed the presence of several rhamnose-rich CWPS biosynthesis loci homologous to those found in *Streptococcus* species. All genes/operons that were targeted for knockout have red highlighting; however, all Rha-CWPS genes except for *gacI* were essential for survival.

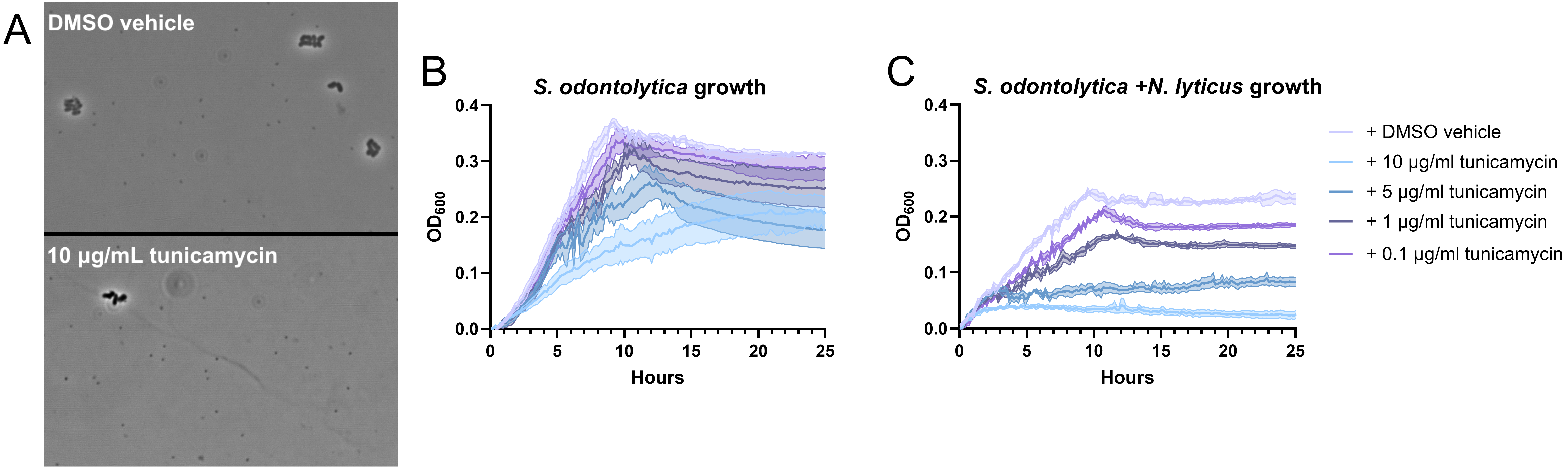

**Supplemental figure 3.** **Optimization of tunicamycin treatment.** Since most Rha-CWPS genes were determined to be essential for survival, a chemical inhibitor of the CWPS synthesis protein TagO called tunicamycin was used to disrupt synthesis. **(A)** Tunicamycin did not appear to inhibit *N. lyticus* cells, as episymbionts were plentiful at all assayed concentrations. **(B)** However, Tunicamycin did inhibit *S. odontolytica* growth at all assayed concentrations, when monitored via growth curve. **(C)** Tunicamycin inhibited *S. odontolytica* synergistically with *N. lyticus* when grown in co-culture, and at 10 μg/ml these combined stressors completely halted growth. Based on these results, tunicamycin treatments were performed at 0.1 μg/ml to minimize host cell stress.

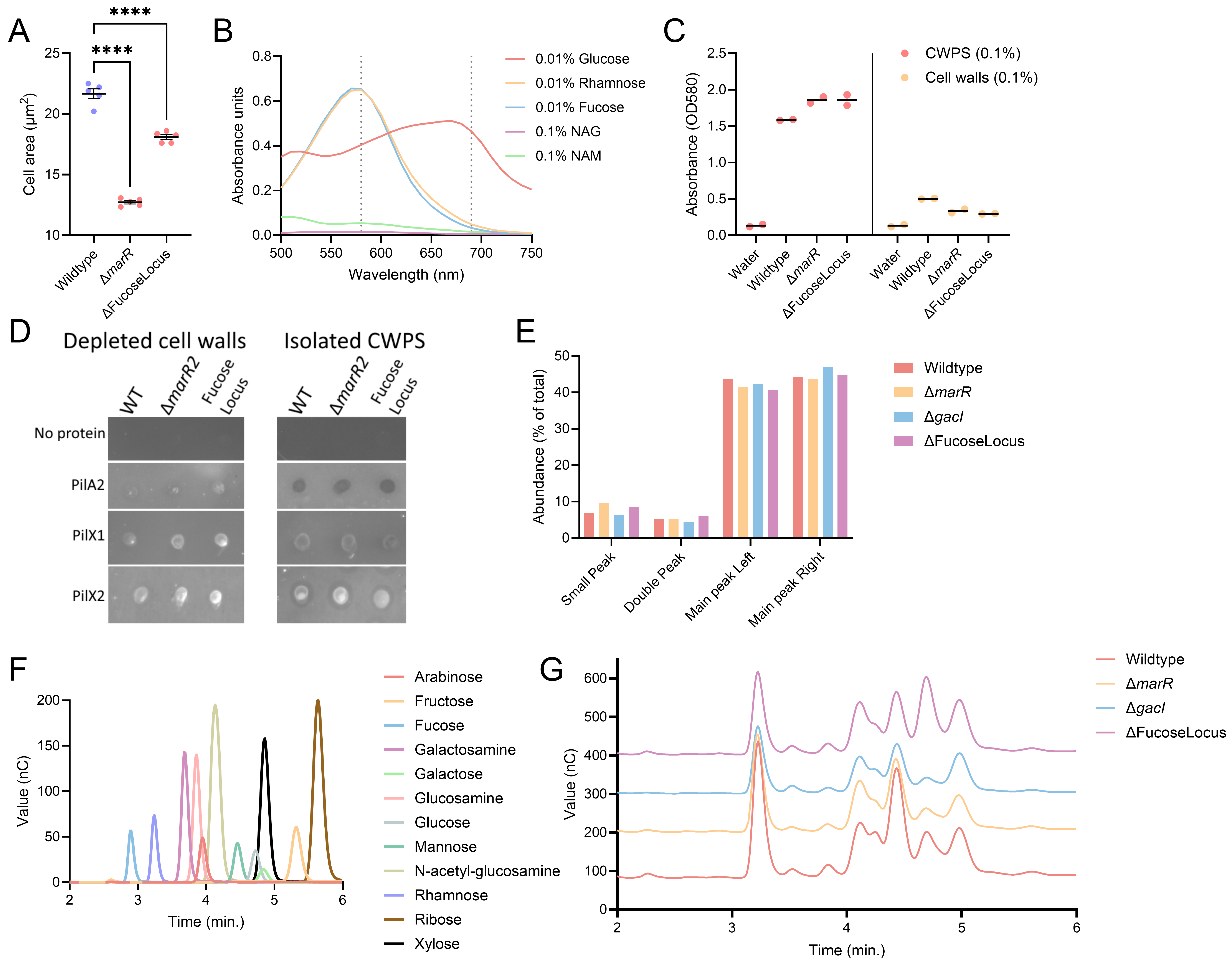

**Supplemental figure 4.** **Analysis of CWPS biosynthesis mutants. (A)** *S. odontolytica* Δ*marR* and *S. odontolytica* ΔFucoseLocus cells were significantly shorter on average than wildtype cells. Since this organism can have pleomorphic shapes, cross-sectional cell area was used instead of cell length. **(B)** The anthrone assay hydrolyzes saccharides into colorimetric products which can be differentiated by absorbance pattern. These monosaccharide standards display how 6-deoxyhexoses generate a product with a relatively narrow absorbance peak around 580 nm, while glucose produces a product with a wider absorbance peak. N-acetylglucosamine (NAG) and N-acetylmuramic acid (NAM) from peptidoglycan contribute little to no product from this reaction. **(C)** The anthrone assay confirms that TCA cleavage liberates the majority of CWPS from cell walls, though some residual saccharides remain attached, particularly in wildtype cells. **(D)** An example replicate of the dot blots prepared with glycan depleted cell walls and the isolated CWPS fractions. **(E)** Using HPLC to separate glycans within collected CWPS fractions, five peaks were detected in all samples. Relative abundances were similar between all treatments. **(F)** When performing HPAEC-PAD, we developed a set of monosaccharide standards common in known Gram-positive CWPS. **(G)** These standards were compared to the peaks present in CWPS hydrolysates to quantify the relative abundance of each component, here shown with representative traces.

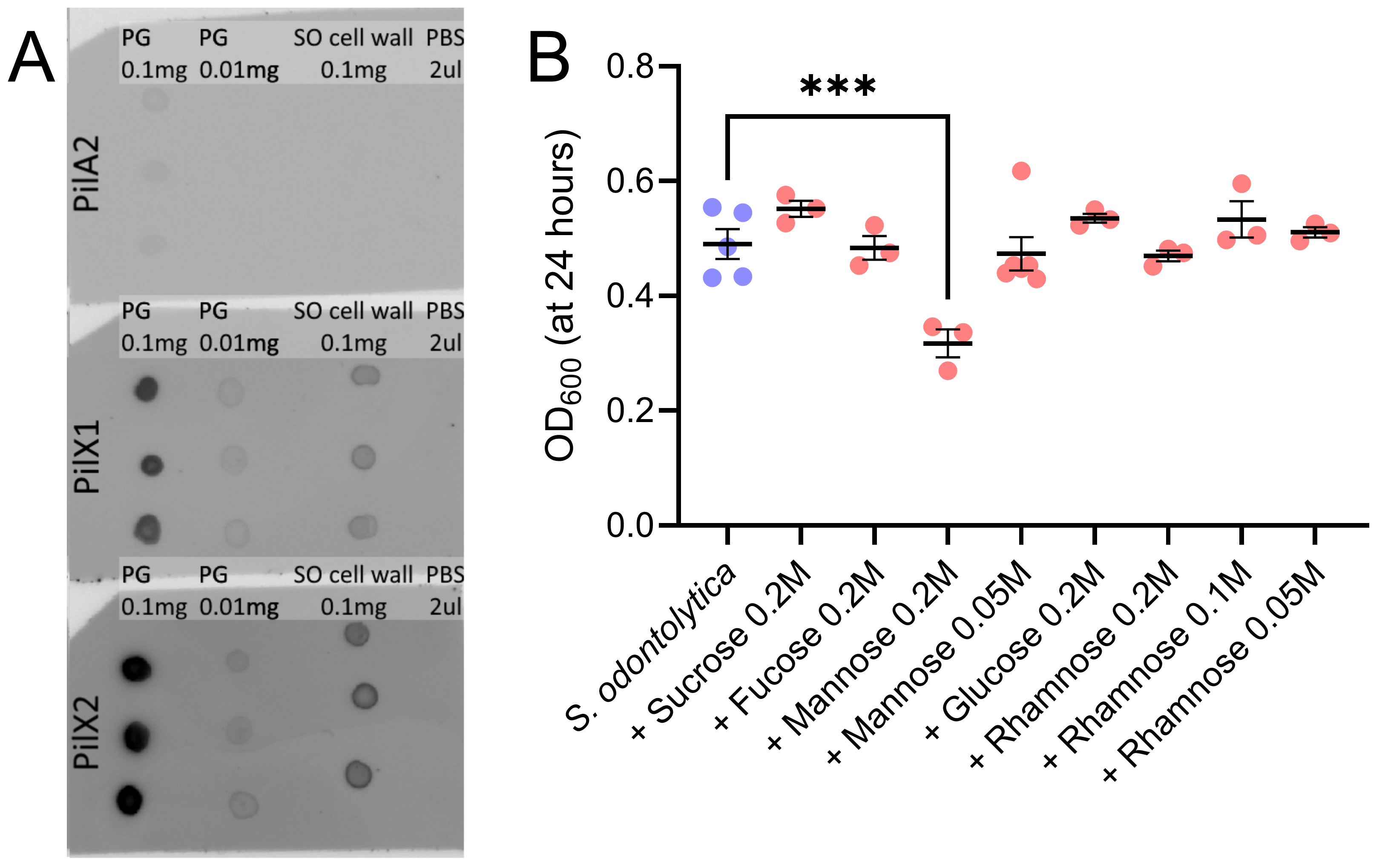

**Supplemental figure 5. Exogenous mannose induces sugar toxicity in *S. odontolytica*.** Cultivation of *S. odontolytica* monocultures within medium supplemented with monosaccharides known to occur in Gram-positive CWPS revealed that there was no solute toxicity at 0.2 M concentrations, however high concentrations of mannose specifically inhibited growth.
